# Human anelloviruses produced by recombinant expression of synthetic genomes

**DOI:** 10.1101/2022.04.28.489885

**Authors:** Dhananjay M. Nawandar, Maitri Trivedi, George Bounoutas, Kevin Lebo, Cato Prince, Colin Scano, Nidhi Agarwal, Erin Ozturk, Jason Yu, Cesar A. Arze, Agamoni Bhattacharyya, Dinesh Verma, Parmi Thakker, Joseph Cabral, Shu-Hao Liou, Kurt Swanson, Harish Swaminathan, Fernando Diaz, Ashley Mackey, Yong Chang, Tuyen Ong, Nathan L. Yozwiak, Roger J. Hajjar, Simon Delagrave

## Abstract

Human anelloviruses are acquired universally in infancy, highly prevalent, abundant in blood, and extremely diverse. Their apparent lack of pathogenicity indicates that they are a major component of the commensal human virome. Despite their being extensively intertwined with human biology, these viruses are poorly understood. A major impediment in studying anelloviruses is the lack of an *in vitro* system for their production and/ or propagation. Here we show that the T cell-derived human cell line MOLT-4 can be transfected with plasmids comprising tandem anellovirus genomes to produce viral particles visualized by electron microscopy. We found that a previously described human anellovirus of the *Betatorquevirus* genus (LY2), as well as a second *Betatorquevirus* detected by sequencing DNA extracted from a human retinal pigmental epithelium (nrVL4619), can be synthesized and produced by these means, enabling further molecular virology studies. Southern blot was used to demonstrate replication, and site-directed mutagenesis of the viral genome was performed to show that the production of anellovirus in this cell line is dependent on the expression of certain viral proteins. Finally, experiments performed in mice using purified nrVL4619 particles produced in MOLT-4 cells demonstrated infectivity *in vivo* in the tissue of origin. These results indicate that anelloviruses can be produced *in vitro* and manipulated to improve our understanding of this viral family which is ubiquitous in humans and many other mammals. Applications of this work to gene therapy and other therapeutic modalities are currently under investigation.

**IMPORTANCE:** Anelloviruses are a major component of the human virome. However, their biology is not well understood mainly due to the lack of an *in vitro* system for anellovirus production and/or propagation. In this study, we used multiple orthogonal measures to show that two different anelloviruses belonging to the *Betatorquevirus* genus can be produced in a T-cell-derived human cell line, MOLT-4, via recombinant expression of synthetic genomes. Additionally, we show that anellovirus particles generated in this *in vitro* system demonstrate infectivity *in vivo*. Our findings enable new molecular virology studies of this highly prevalent, non-pathogenic, and weakly immunogenic family of viruses, potentially leading to therapeutic applications.

## INTRODUCTION

Anelloviruses are small non-enveloped viruses encoding a circular single-stranded, negative-sense DNA genome. Human anelloviruses can be classified into three genera - *Alphatorquevirus* (torque teno virus [TTV]), *Betatorquevirus* (torque teno mini virus [TTMV]), and *Gammatorquevirus* (torque teno midi virus [TTMDV]). Their genome sizes are approximately 3.9 kilobases (kb) for *Alphatorqueviruses*, 2.9 kb for *Betatorqueviruses*, and 3.2 kb for *Gammatorqueviruses*. Each contains a non-coding region with an ∼100-base pair, guanine-cytosine (GC)-rich sequence and several overlapping open reading frames (ORFs), the largest of which, ORF1 (∼700-800 aa), is the putative viral capsid protein^1^. Genome replication is thought to occur via a rolling circle mechanism common to other circular DNA viruses and employs host polymerases.

Since their discovery in 1997^2^, anelloviruses have been found in many biological samples including blood, nasal secretions, saliva, bile, feces, tears, semen, breastmilk, and urine, suggesting broad tropism for different cell and tissue types^3–6^. A study of 44 healthy infants showed that they all acquired anelloviruses within their first year of life^7^. Once acquired, anelloviruses appear to persist in the body and avoid clearance by the immune system; the detection of the same type of anellovirus in samples from an individual dialysis patient collected 16 years apart supports the theory that people can remain life-long anellovirus carriers^8^.

Human anelloviruses have not been convincingly associated with any disease^9^. In fact, potentially beneficial effects on human health have been suggested for anelloviruses^4^. For instance, acquisition of anelloviruses by newborns^7,10^ could promote the development and maturation of the immune system^11^. Such results are concordant with a long history of co-evolution between the virus and the host, eventually leading to commensal or even mutualistic relationships^12^.

Despite their discovery more than 25 years ago and their very high prevalence in the human population, relatively little is known about the biology of anelloviruses because of the lack of an *in vitro* cell culture system and other tools such as reliable serologic assays and animal models^12,13^. In this study, we demonstrate that a human lymphoblastic cell line, MOLT-4, may be employed for the production of two distinct anelloviruses belonging to the *Betatorquevirus* genus. In addition to characterizing viral gene expression and replication, we provide the first evidence to our knowledge of the production of purified virus particles for analysis by transmission electron microscopy (TEM) as well as infectivity *in vivo*. The *in vitro* production model and related tools described herein promise to advance the study of previously neglected human anelloviruses, including their molecular virology and virus-host interactions.

## MATERIALS AND METHODS

### Dissection of human samples

Human eyes were obtained through the National Disease Research Institute (NDRI) and were dissected within 24-48 hours of procurement. Each individual eye was placed on a dissecting plate, and the sclera was incised at a point between the cornea and the optic nerve using a razor blade. From that point, the sclera was cut all the way around. The aqueous humor and vitreous humor were isolated separately. The choroidal layer was then removed, and the retina slowly peeled off and processed. Other compartments in the eye that were isolated and analyzed were the sclera, the iris, the cornea, the conjunctiva, and the optic nerve.

### DNA extraction and processing

Dissected tissue sections were homogenized and DNA was extracted with a PureLink viral DNA/RNA kit from Invitrogen (catalog # 12280050). The samples were processed essentially according to the manufacturer’s protocol. The extracted DNA then underwent rolling circle amplification (RCA) following the procedure outlined by Arze *et al* for a final volume of 20 µL^14^. The presence of *Anelloviridae* in the samples was tested by PCR with pan-anellovirus primers developed by Ninomiya et al^7^.

### Illumina library preparation and sequencing

Post-RCA DNA was diluted to a volume of 50 µL to reduce viscosity of the samples and then the concentration of DNA was assessed by Qubit 4 Fluorometer (Thermo Fisher Scientific). Post-RCA DNA was library-prepped using the Nextera DNA kit (Illumina). The samples were prepared following the manufacturer’s protocol for 100-500 ng input. Post-RCA DNA was also library-prepped using SureSelect XT HS2 DNA Reagent Kit (Agilent) with target enrichment RNA probes. These target enrichment RNA probes were specifically designed to tile across *Anelloviridae* sequences from our database and were biotinylated to enable capture using streptavidin beads. Library quality control was carried out with D5000 ScreenTape on a 4200 TapeStation (Agilent). All libraries were then sequenced on either an iSeq 100 or a NextSeq 550 (Illumina).

### Nanopore library preparation and sequencing

Post-RCA DNA was debranched and fragmented to 20 kb-sized fragments following the NanoAmpli-Seq protocol^15^. 4.5 μg of RCA material was diluted in 65 μL of nuclease-free water and treated with 2 μL of T7 endonuclease I (New England Biolabs) for 5 minutes at room temperature. The reaction was then loaded in a g-TUBE (Covaris) and centrifuged at 1800 rpm (304 relative centrifugal force [RCF]) for 4 minutes. The g-TUBE was then reversed, and the centrifugation process was repeated. An additional round of T7 endonuclease I and g-TUBE was performed before the mixture was then cleaned up with SPRI beads at a ratio of 1.8 × with a final elution in 20 μL of nuclease-free water. The concentration of DNA was assessed by Qubit 4 fluorometer (Thermo Fisher Scientific). The fragmented samples were then library-prepped with a SQK-LSK109 kit (Oxford Nanopore Technologies) following the manufacturer’s protocol. Additionally, post-RCA DNA was debranched and fragmented to 6-8 kb-sized fragments using the above-mentioned protocol with the modification of the g-TUBE (Covaris) being centrifuged at 13200 rpm (16363 RCF) for 30 seconds. The samples were prepared with the SureSelect XT HS2 DNA Reagent Kit (Agilent) with biotinylated target enrichment RNA probes specifically designed for *Anelloviridae* following the manufacturer’s protocol with an increased elongation to 6 minutes in amplification steps. The samples were then library-prepped with the SQK-LSK109 kit (Oxford Nanopore Technologies) following the manufacturer’s protocol. Libraries were loaded onto a R9.5 (FLO-MIN107) flow cell and placed onto the MinION Mk1B (Oxford Nanopore Technologies) and run for 48 hours. Only flow cells that passed the manufacturer’s flow cell check test were used.

### Sequence quality control

Both Illumina and Nanopore raw sequencing reads were subjected to quality control utilizing FastQC on the sequence datasets derived from each instrument^16^. Reports generated by FastQC for each individual sample were then aggregated into a single report using the MultiQC^17^ utility. Metrics from these reports influenced parameter selection to downstream quality control steps during analysis.

Illumina sequence data were filtered to remove low-quality sequences and common adapters using bbduk with the following parameters: *ktrim=r, k=23, mink=11, tpe=t, tbo=t, qtrim=rl, trimq=20, minlength=50, maxns=2*^18^. The target contaminant file used was assembled by pulling contaminant sequences from NCBI GenBank covering several bacterial and human genetic elements and common laboratory synthetic sequences to be removed.

Nanopore sequence data were filtered to remove adapter sequences with porechop using default parameters followed by quality and length filtering using filtlong with parameters *--min_length 2000 --keep_percent 90* ^19,20^ Reads passing quality control were mapped to anellovirus contig sequences with the following parameters: -cx map-ont. The resultant PAF file was both visualized in Alvis and parsed to identify best hits to the reference contig sequences, and these reads were further analyzed with pairwise alignments in Geneious (Biomatters) with the MAFFT alignment plug-in with the G-INS-i algorithm^21^. These long reads were used to validate the assembled short-reads and to verify that these contigs were not chimeras.

Next, human sequences were removed in two passes with both NextGenMap and BWA against the GRCh37/hg19 build of the human reference genome^22–24^. NextGenMap was run with parameters *--affine, -s 0*.*7*, and *-p*, and BWA was run with default parameters. Mapped reads output in SAM file format were converted to paired-end FASTQ format with both SAMtools and Picard’s SamToFastq utility configured with the parameter *VALIDATION_STRINGENCY=“silent”*^24,25^.

rRNA contaminants and common laboratory bacterial contaminants were removed with bbmap with the following parameters: *minid=0*.*95, bwr=0*.*16, bw=12, quickmatch=t, fast=t, minhits=2*. An accounting of all reference sequences screened against can be found in the provided supplementary data^18^.

Finally, we de-duplicated the short-read data passing all QC and decontamination steps to speed up and aid in genome assembly quality by using clumpify configured with the parameter *dedupe=t*^18^.

### Genome assembly

Short, trimmed, decontaminated, and de-duplicated sequencing reads were assembled with metaSPAdes, with the error correction module disabled via the use of the *--only-assembler* parameter. The resulting contigs were filtered with PRINSEQ lite using the parameters *out_format 1, -lc_method dust*, and *lc_threshold 20* ^26,27^. Contigs passing this filtering step were then clustered at 99.5% similarity to remove any duplicate sequences via the VSEARCH software’s *cluster_fast* algorithm using default parameters. Any putative complete, circular genomes were recovered from contigs using ccfind, with all parameters set to defaults^28,29^.

### Long-read error correction

Nanopore reads classified as anellovirus sequences were error-corrected using paired short-read data utilizing racon^30^. First, short reads classified as anellovirus were mapped to long anellovirus reads using BWA’s *mem* algorithm with default parameters^23^. The resulting SAM alignment and the short reads and long reads used to produce the alignment were supplied to racon for error correction^30^. Execution of racon was conducted using default parameters for three rounds of error correction until the polished product showed no variation from the previous iteration.

### Anellovirus contig identification

Assembled contigs were screened using NCBI’s blastn software, with default parameters, to identify putative anellovirus sequences using a custom in-house anellovirus database consisting of 728 curated sequences^31^.

### Anellovirus genome annotation

ORF sequences were identified and extracted from assembled anellovirus contigs using the OrfM software with parameters configured to print stop codons (*-p*) as well as ORFs in the same frame as a stop codon (*-s*) and constrained to ORF sequences longer than 50 amino acids (*-m 150*)^32^.

Predicted ORF sequences were further filtered using seqkit’s *seq* and *grep* utilities to subdivide ORF sequences into bins corresponding to ORF1, ORF2, and ORF3^33^. ORF1 sequences were identified by filtering ORF sequences using seqkit *seq* for those no shorter than 600 amino acids (*-m 600)* and using seqkit *grep* to search through the ORF sequence data (*-s)* for the conserved motif YNP*X*^*2*^D*X*G*X*^*2*^N with a regular expression (*-r*)-based pattern (*-p “YNP*.*{2}D*.*G*.*{2}”*) ^34^. Similarly, ORF2 sequences were recovered using the conserved motif W*X*^*7*^H*X*^*3*^C*X*C*X*^*5*^H previously identified in literature through seqkit’s *grep* utility (*-p “W*.*{7}H*.*{3}C*.*C*.*{5}H”*) ^35^.

ORF3 sequences were predicted by utilizing the presence and coordinate positions of predicted ORF1 and ORF2 sequences on the same contig. Predicted ORF3s use a stop codon downstream of those used by ORF1 and their reading frames are different from those of ORF1 and ORF2 sequences. Additionally, parsing the ORF3 sequences from internal datasets (median length: 68 aa, minimum length: 50 aa, maximum length: 159 aa through MEME revealed the presence of two previously unknown and highly conserved motifs located near the 3’ end of ORF3^36^. Both novel motifs were also utilized to identify ORF3 sequences using seqkit’s *grep* command.

Identified ORF sequences required an additional trimming step as OrfM produces ORF calls with peptides upstream of canonical start codons. ORF1 sequences were timed to the proper start codon via an in-house written python script that used the presence of the arginine-rich region to identify the first methionine located upstream of it in the direction of the 5’ end. In some cases, a non-canonical start codon was predicted as the ORF1 start codon by searching for a threonine-proline-tryptophan or threonine-alanine-tryptophan tripeptide directly upstream of the arginine-rich region. ORF2 and ORF3 sequences were trimmed to the first start codon identified nearest the 5’ end of the sequence.

### Anellovirus genera classification

Anellovirus contig sequences were identified into one of the three known genera by use of the tblastx software to conduct a homology search against a custom in-house database consisting of 720 curated and classified anellovirus sequences^31^. The top hits that contained suitable coverage across the majority of the contig sequence were then used in genera classification.

### Primer walking and genome recovery

Primers were designed around regions of inconsistencies between the long-read and short-read sequencing data. Post-RCA DNA was amplified using these primers with a Q5 Hot Start polymerase (New England Biolabs). The product was run on a 2% gel to confirm specific binding before sending the PCR product to GeneWiz for Sanger sequencing. Sanger sequencing results were analyzed using Geneious bioinformatics software (Biomatters).

### Plasmid construction

#### Plasmid containing a single copy of WT LY2, all ORF1 KO LY2 and all ORF2 KO LY2

The sequence of a previously described anellovirus LY2 (GenBank accession number: JX134045.1) that belongs to the *Betatorquevirus* genus was synthesized by Integrated DNA Technologies into pUCIDT-Kan plasmid (pUCIDT-LY2)^37^. Esp3I restriction cut sites were added on each side of the genome in this plasmid to enable subcloning, scarless restriction digestion, and ligation of the two ends of the genome to make double-stranded circular genomes. The template plasmid was amplified with the following primers: FWD 5’-ACAGCTCTTCAAGGCGTCTCACCTAATAAATATTCAACAGGAAAACCACCTAATTTAAATTGCC-3’ and REV 5’-ACAGCTCTTCAGTGCGTCTCATAGGGGGTGTAAGGGGGCGTAG-3’. PCR reactions (50 µl) contained 1.0-unit Phusion DNA polymerase (New England Biolabs catalog # M0419), 1X Phusion HF buffer, 200 µM dNTPs (New England Biolabs # N044), 0.5 µM of each primer (synthesized at Integrated DNA Technologies), 3% DMSO, and 1 ng of template DNA. All PCR reactions were run with the following parameters: initial denaturing at 98°C for 30 seconds followed by 40 cycles of denaturing at 98°C for 15 seconds, annealing at 60°C for 30 seconds, extension at 72°C for 3 minutes, and a final extension at 72°C for 10 minutes.

Purified PCR product was cloned into a pcDNA 6.2/V5-PL-DEST (Thermo Fisher Scientific catalog # 12537162) destination plasmid in a one-pot reaction containing 50 ng of destination vector, 30 ng of PCR product, 1 × BSA, 1 × T4 DNA ligase buffer (New England Biolabs), 10 units of BspQI (New England Biolabs catalog # R0712)), and 400 units of T4 DNA ligase (New England Biolabs catalog # M0202). Cloning reaction was incubated at 50°C for one hour followed by 15 minutes at 16°C.

Constructs to knock out the protein expression of either all ORF1 variants (all ORF1 KO LY2) or all ORF2 variants (all ORF2 KO) were designed by inserting a premature stop codon – Cysteine9-STOP and Arginine13-STOP, respectively. These mutations and the surrounding sequences were ordered as gBlocks (Integrated DNA Technologies) for restriction digest cloning into WT LY2. The gBlocks and WT LY2 plasmid were digested with SpeI-HF and SalI-HF (New England Biolands). The plasmid was further treated with QuickCIP (New England Biolands). The digested products were separated by gel electrophoresis, purified and then ligated. All clones were verified through Sanger sequencing at Genewiz.

#### Plasmid containing two copies of LY2 in tandem

A plasmid harboring two copies of the LY2 genome arranged in a tandem configuration was assembled using a Golden Gate cloning method. The LY2 genome was subcloned into Level 1 plasmids as genome 1 (G1) and genome 2 (G2) with PCR primers containing different Esp3I overhangs for later assembly. The plasmids were amplified by PCR with forward G1-F 5’-ACAGCTCTTCAAGGCGTCTCAATGGTAATAAATATTCAACAGGAAAACCACCTAATTTAAATTGCC-3’ and reverse G1-R 5’-ACAGCTCTTCAGTGCGTCTCATAGGGGGTGTAAGGGGGCGTAG-3’ for G1; and forward G2-F 5’-ACAGCTCTTCAAGGCGTCTCACCTAATAAATATTCAACAGGAAAACCACCTAATTTAAATTGCC-3’ and reverse G2-R 5’-ACAGCTCTTCAGTGCGTCTCATTCAGGGGGTGTAAGGGGGCGTAG-3’ for G2. PCR reactions (50 µl) contained 1.0 unit of Phusion DNA polymerase (New England Biolabs catalog # M0419), 1 × Phusion HF buffer, 200 µM of dNTPs (New England Biolabs # N044), 0.5 µM of each primer (synthesized at Integrated DNA Technologies), 3% DMSO, and 1 ng of template DNA. All PCR reactions were run with the following parameters: initial denaturing at 98°C for 30 seconds followed by 40 cycles of denaturing at 98°C for 15 seconds, annealing at 60°C for 30 seconds, extension at 72°C for 3 minutes, and a final extension at 72°C for 10 minutes. For assembling the tandem genome plasmid, the destination plasmid, G1 subclone, and G2 subclone were cloned in a one-pot Golden Gate reaction containing 50 ng of the destination plasmid, 30 ng of each genome subclone, 1 × BSA, 1 × T4 DNA ligase buffer, 10 units of Esp3I (New England Biolabs catalog # R0734), and 400 units of T4 DNA ligase (New England Biolabs catalog # M0202). The cloning reaction was run at 37°C for 15 minutes, 20 cycles at 37°C for 2 minutes followed by 15°C for 5 minutes, at 37°C for 15 minutes, at 50°C for 5 minutes, and at 80°C for 5 minutes.

#### Plasmid containing two copies of nrVL4619 in tandem

A single copy of the nrVL4619 genome, flanked by BsaI cut sites, was synthesized by GenScript into a pUC57-Kan vector. The nrVL4619 genome was excised and separated from its plasmid backbone using BsaI-HFv2 (New England Biolabs catalog # R3733) and PvuI-HF restriction enzymes (New England Biolabs catalog # R3150); the excised band was purified and ligated to itself to form an *in vitro* circularized (IVC) genome. A plasmid containing tandem copies of nrVL4619 was cloned by linearizing both the IVC genome and a plasmid containing a single copy of nrVL4619 (described above) with NheI-HF restriction enzyme and ligating with T4 DNA ligase (New England Biolabs). All clones were verified through Sanger sequencing at Genewiz.

### In vitro circularization (IVC) of LY2 genome

A circularizable LY2 plasmid was digested with Esp3I (New England Biolabs catalog # R0734) and PvuI-HF (New England Biolabs catalog # R3150), separating the genome and plasmid backbone by gel electrophoresis. To purify the LY2 genome from the excised gel, the gel was placed in 10K MWCO SnakeSkin dialysis tubing (Thermo Fisher Scientific catalog # 88242), electroeluting in a 1 × TAE gel box for 18 hours at 40V. Buffer/DNA solution was removed from tubing, incubating mixture with concentrated (2,000,000 units/mL) T4 DNA ligase and ligase buffer (New England Biolabs) at 16°C for 12 hours. Ligated solution was concentrated in 30kD Microsep Advance centrifuge tubes (Pall Corporation).

### Cell culture

MOLT-4 cells were obtained from the National Cancer Institute. Cells were scaled-up and maintained in suspension culture in complete growth medium (Gibco’s RPMI 1640 with 10% fetal bovine serum [FBS], supplemented with 1 mM sodium pyruvate, Pluronic F-68 [0.1%], and 2 mM L-glutamine) at 37°C with 5% CO_2_. Cells were seeded into shake flasks (2-L, flat-bottomed, Erlenmeyer flask), each with a working volume of 800 mL, at a density of 0.1E+06 viable cells/mL and cultured in an orbital shaker (New Brunswick Innova 2100, 19-mm circular orbit) at 37°C and 100 rpm with >85% relative humidity (RH) for 4 days.

### Transfection of MOLT-4 cells

MOLT-4 cells were transfected with the indicated plasmids either by nucleofection or electroporation.

For nucleofection at 25 mL scale, cells were counted using the BioProfile FLEX2 analyzer (Nova Biomedical), and 10^7^ cells were pelleted by spinning at 200 × g for 10 minutes. Pelleted cells were resuspended in SF Cell Line Nucleofector Solution with added supplement (Lonza catalog # V4XC2024). 25 µg of the plasmid to be transfected (Aldevron) was added to the resuspended cells and nucleofected using the CM-150 program on the 4D-Nucleofector X Unit (Lonza). Nucleofected cells were allowed to recover in a 37°C incubator with 5% CO_2_ for 20 minutes, after which they were added to a flask containing pre-warmed complete growth medium.

For electroporation at 25 mL scale, 10^7^ pelleted cells were resuspended in homemade 2S Chica buffer (5 mM KCl, 15 mM MgCl_2_, 15 mM HEPES buffer solution, 150 mM Na_2_HPO_4_ pH 7.2, 50 mM sodium succinate). 100 µg of the plasmid to be transfected (Aldevron) was added to the resuspended cells and electroporated using a NEPA21 electroporator (Bulldog Bio). Electroporated cells were then transferred to a flask containing pre-warmed complete growth medium.

Transfected cells were allowed to incubate at 37°C with 5% CO_2_ and harvested at the indicated times.

### Western blotting

Cell pellets were resuspended in lysis buffer containing 50 mM Tris pH 8.0, 0.5% Triton-X100, 100 mM NaCl, and 1 × Halt protease inhibitor cocktail (Thermo Fisher Scientific catalog # 78439), followed by two rounds of freeze-thawing. The cell lysates were clarified by centrifugation at 10,000 × g for 30 minutes at 4°C, and the protein concentration was quantified using Pierce BCA Protein Assay Kit (Thermo Fisher Scientific catalog # 23227) according to the manufacturer’s protocol. Equal amounts of the cell lysates were mixed with loading dye and Bolt sample reducing agent (Thermo Fisher Scientific catalog # B0009), followed by boiling at 95°C for 5 minutes.

For ORF2 and GAPDH, proteins were separated on Bolt 4-12% Bis-Tris gel in 1 × Bolt MOPS SDS running buffer (Thermo Fisher Scientific catalog # B0001). Separated proteins were electro-transferred to nitrocellulose membrane using Trans-Blot Turbo Transfer System (Bio-Rad). For ORF1, proteins were separated on Bolt 12% NU-PAGE gel and transferred to nitrocellulose membrane at 100 volts for 1.5 hours by a wet transfer method using cold 1 × Bolt transfer buffer (Thermo Fisher Scientific catalog # BT0006) supplemented with 20% methanol.

After transfer, membranes were blocked in Odyssey blocking buffer (LI-COR) for 1 hour and then incubated with relevant primary antibodies overnight. Anti-ORF2 antibody was generated by immunizing rabbits with purified full-length ORF2 protein expressed in *E. coli*. Anti-ORF1 antibody was generated in mice against the jelly roll domain of the ORF1 protein. Anti-ORF2 and -ORF1 antibodies were used at a concentration of 1:500. Anti-GAPDH antibody (Cell Signaling Technologies, catalog # 97166) was used at a concentration of 1:1000 to detect GAPDH as a loading control.

Membranes were washed three times by rocking in a mixture of tris-buffered saline (TBS) and Polysorbate 20 for 10 minutes each. Membranes were then incubated in the relevant secondary antibodies conjugated with fluorescent dyes. Secondary antibodies used were goat anti-mouse IgG paraproteins (IRDye 680RD, LI-COR, catalog # 926-68070, 1:5000 dilution) and goat anti-rabbit IgG (IRDye 680RD, LI-COR, catalog # 926-68071, 1:5000 dilution). Specific immunoreactive proteins were detected using Odyssey DLx imaging system (LI-COR).

### Reverse transcriptase quantitative PCR (RT-qPCR)

Transfected MOLT-4 cells were harvested by centrifugation at 500 × g for 5 minutes. Pelleted cells were lysed using 700 µl QIAzol lysis reagent (Qiagen catalog # 79306), followed by RNA extraction using miRNeasy Mini Kit (Qiagen, catalog # 217004) as per the manufacturer’s protocol. Additional DNAse treatment was also performed on the harvested RNA using RQ1 RNase-Free DNase (Promega, catalog # M6101) according to the manufacturer’s protocol to remove any carryover of double-stranded or single-stranded DNA. cDNA synthesis was performed from DNAse-treated RNA with oligo(dT) primer using SuperScript III First-Strand Synthesis System (Invitrogen, 18080-051). qPCR was performed in triplicate using gene-specific primers with SYBR Green PCR Master Mix (Thermo Fisher Scientific catalog # 4309155) in QuantStudio 5 Real-Time PCR machine (Applied Biosystems). Relative quantity was calculated using human GAPDH as a loading control. Gene-specific primers with the following sequences were synthesized at Integrated DNA Technologies: For LY2:F: CTTATTACTACAGAAGAAGACGGTAC and R: AAAGGGCGTCTAATCCAACC. For GAPDH, F: ACCACAGTCCATGCCATCAC and R: TCCACCACCCTGTTGCTGTA.

### Southern blotting

Isolation of total DNA from a total of 10^7^ transfected MOLT-4 cells was done using DNeasy Blood & Tissue Kit (Qiagen catalog # 69504). Isolated DNA was digested with either NcoI-HF (New England Biolabs # R0193) or NcoI-HF and DpnI restriction enzymes (New England Biolabs catalog # R0176) overnight at 37°C. NcoI-HF cuts the LY2 genome once. The digested samples were separated by gel electrophoresis and subsequently transferred overnight onto a Hybond-N+ membrane. The membrane was hybridized overnight in ULTRAhyb hybridization buffer (Thermo Fisher Scientific catalog # AM8670) and probed using in-house-generated, biotin-labeled oligos to detect the LY2 genome. These LY2-specific probes were made by random priming and labeled with biotin using the BioPrime Array CGH Genomic Labeling System (Invitrogen catalog # 18095012). Membranes were incubated with IRDye800 and imaged using Odyssey DLx imaging system (LI-COR).

### Isopycnic centrifugation

#### Cesium chloride (CsCl) linear gradients

Four days after transfection, MOLT-4 cells were harvested by centrifugation, followed by resuspension in lysis buffer containing 50 mM Tris pH 8.0, 0.5% Triton-X100, 100 mM NaCl, and 1 × Halt protease inhibitor cocktail (Thermo Fisher Scientific catalog # 78439). Resuspended cells underwent by two rounds of freeze-thawing and addition of equal volumes of buffer containing 50 mM Tris pH 8.0 and 2 mM MgC_l2_. Cell lysates were subjected to treatment with 100 U/mL of Benzonase endonuclease (Sigma-Aldrich catalog # E8263) and nutation at room temperature (RT) for 90 minutes. Benzonase-treated cell lysates were clarified at 10,000 × g for 30 minutes at 4°C to pellet any cellular debris.

CsCl linear gradients were prepared by overlaying 8.5 mL of 1.46 g/cm^3^ CsCl solution with 8.5 mL of 1.2 g/cm^3^ CsCl solution in 17 mL Ultra-Clear tubes (Beckman Coulter), which were then spun at a 45-degree angle and a speed of 20 rpm for 13.5 minutes using Gradient Master (BioComp).

2 mL of CsCl solution from the top of the tube was replaced with 2 mL of the processed MOLT-4 cell lysates. The sample-containing tube was spun at 31,000 RPM for 18 hours using SW 32.1 rotor (Beckman Coulter). 1-mL fractions were collected from the bottom of the tube. The refractive index of each fraction was measured using Refracto handheld refractometer (Mettler Toledo) to calculate density. Each fraction was desalted using a desalting kit (Thermo Fisher Scientific catalog # 89851) and then subjected to DNAse-protected qPCR assay as described below.

#### Iodixanol linear gradients

MOLT-4 cells were harvested and processed as described above for CsCl linear gradients. To prepare iodixanol linear gradients, 13 mL of 60% OptiPrep (Sigma-Aldrich catalog # D1556) was overlayed with 13 mL of 20% OptiPrep in 26.3-mL polycarbonate tubes, which were then spun at a 46-degree angle and a speed of 20 rpm for 16 minutes using Gradient Master (BioComp). The sample-containing tube was spun at 347,000 × g and 20°C for 3 hours using Type 70 Ti rotor (Beckman Coulter). 1-mL fractions were collected from the top of the tube. The refractive index of each fraction was measured using Refracto handheld refractometer (Mettler Toledo) to calculate density. Each fraction was then subjected to DNase-protected qPCR assay as described below.

### DNase-protected qPCR assay

5 µl of the sample to be titrated was incubated with 200 U of DNAse I endonuclease (Thermo Fisher Scientific catalog # 18047019) in a 20-µl reaction. The reaction was incubated at 37°C for 2 hours, followed by inactivation of DNase I at 95°C for 10 minutes. 4 µl of the 1:10 diluted DNase reaction was subjected to qPCR analysis in a 20-µl reaction using TaqMan Universal PCR Master Mix (Thermo Fisher Scientific catalog # 4304437) according to the manufacturer’s protocol. Primer and probe sequences are listed in Table 1.

**TABLE 1.**
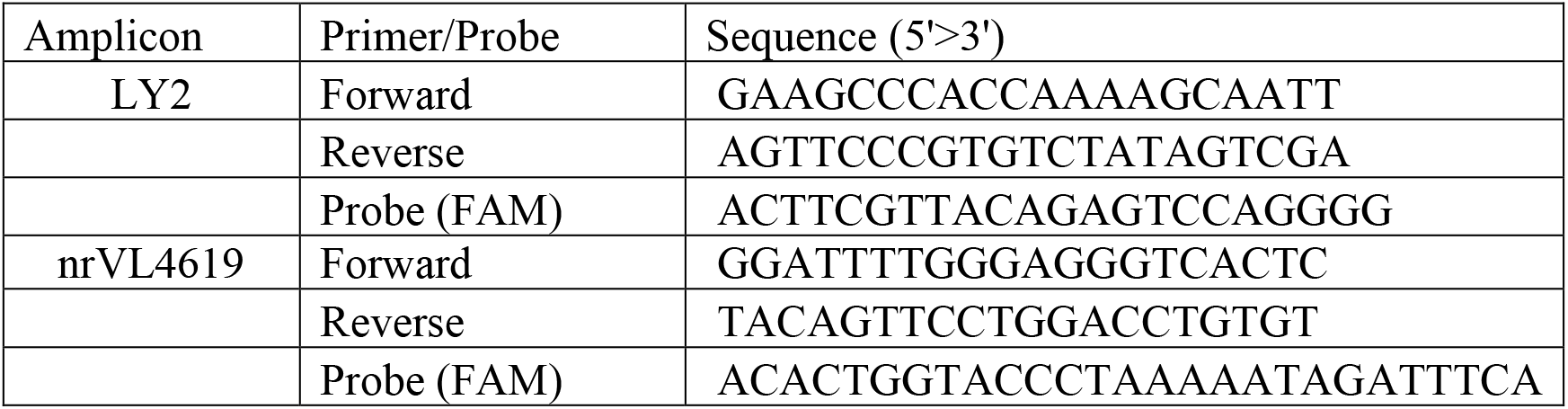
Primer and probes designed to quantify LY2 and nrVL4619 titers in virus preparations produced from MOLT-4 cells.

### LY2 scale-up production

#### Nucleofection

Cells were counted using BioProfile FLEX2 analyzer (Nova Biomedical), and 2×10^9^ viable cells were pelleted using Sorvall BIOS A floor model centrifuge (Thermo Fisher Scientific) in 1-L bottles at 500 RCF for 30 minutes. The supernatant was discarded, the pellets were resuspended in 20 mL of P3 solution with added supplement (Lonza), and 2 mg of the plasmid encoding tandem copies of the LY2 genome (Aldevron) was added. The cells were nucleofected using 4D Nucleofector LV Unit (Lonza) and collected in 5 mL of complete growth medium. The nucleofected cells were then transferred to 600 mL of pre-warmed complete growth medium in a shake flask and incubated in a shaker at 37°C and 100 rpm with 5% CO_2_ and >85% RH for 1 hour.

After incubation, the cells were counted using BioProfile FLEX2 analyzer (Nova Biomedical). They were then diluted to 4×10^5^ viable cells/mL in pre-warmed complete growth medium in shake flasks (800 mL maximum working volume) and incubated in a shaker at 37°C and 100 rpm with 5% CO_2_ and >85% RH for 4 days.

#### Harvest and cell lysis

Four days after nucleofection, cells were counted using BioProfile FLEX2 analyzer (Nova Biomedical). Cells were then harvested by pelleting using Sorvall BIOS A floor model centrifuge (Thermo Fisher Scientific) at 1000 RCF for 30 minutes, and supernatant was discarded. Cell pellets were resuspended in 30 mL of 20 mM Tris pH 8, 100 mM NaCl, and 2 mM MgCl_2_buffer, lysed using LM10 Microfluidizer (Microfluidics) at 10,000 psi, and washed with 30 mL of the same buffer to make a final cell lysate volume of 60 mL. Then the cell lysates were treated with 1 × Halt protease inhibitor cocktail (Thermo Fisher Scientific) and 100 U/mL Benzonase endonuclease (Sigma-Aldrich) and incubated for 1.5 hours on a stir plate at RT. Next, 0.5% Triton X-100 detergent was added to the cell lysates and returned to incubate at RT on the stir plate for 45 minutes. The treated cell lysates were then centrifuged using 5810 R benchtop centrifuge (Eppendorf) at 10,000 RCF for 30 minutes at 4°C to pellet any cellular debris. Cellular debris was discarded, and the supernatant (lysate) was purified using density gradients.

#### CsCl step gradient

A CsCl step gradient was prepared by underlaying 30 mL benzonase treated and clarified lysate with 3 mL 1.2 g/L CsCl solution and 3 mL 1.4 g/L CsCl solution made in 30 mM Tris and 100 mM NaCl (TN) buffer in 38.6 mL Ultra-Clear ultracentrifuge tubes (Beckman Coulter). Next, the tubes were ultracentrifuged using Optima XE (Beckman Coulter) at 31,000rpm and 10°C for 3 hours. After the spin, the band at the junction of the 1.2 g/L and 1.4 g/L CsCl was extracted and transferred to 3-12 mL Slide-A-Lyzer dialysis cassettes with a molecular weight cutoff (MWCO) of 10K (Thermo Fisher Scientific). The membranes were placed in 1 × Dulbecco’s phosphate-buffered saline (DPBS) with Mg and Ca salts (Gibco), 0.001% Pluronic F-68 (Gibco), and 100 mM NaCl as a dialysis buffer overnight (O/N) on a stir plate at 4°C.

#### CsCl linear gradient and concentration

A CsCl linear gradient was prepared by underlaying 15 mL 1.2 g/L CsCl solution and 15 mL 1.4 g/L CsCl solution in a 30 mL OptiSeal ultracentrifuge tube (Backman Coulter) and spinning using Gradient Master 108 (BioComp) at a 45-degree angle and a speed of 20 RPM for 13.5 minutes. Next, the top 3 mL of CsCl solution was replaced by 3 mL of dialyzed step gradient fraction. The tubes were then ultracentrifuged at 25,000 rpm and 10°C for 18 hours. After the O/N spin, 1 mL fractions were collected in 96 mL-deep well plates from the bottoms of the tubes. Refractive index of each fraction was measured using Refracto handheld refractometer (Mettler Toledo) to calculate density. An aliquot of each fraction was desalted using Zeba 96-well spin desalting plates (Thermo Fisher Scientific) to remove any CsCl and analyzed for LY2 titer using DNAse qPCR. Fractions of interest were determined based on qPCR titer and density. They were then pooled and transferred to 3-12 mL Slide-A-Lyzer dialysis cassettes with a MWCO of 10K (Thermo Fisher Scientific). The membranes were placed in 1 × DPBS with Mg and Ca salts (Gibco), 0.001% Pluronic F-68 (Gibco), and 100 mM NaCl as a dialysis buffer O/N on a stir plate at 4°C. The dialyzed sample was concentrated ten-fold using Amicon Ultra centrifugal filter units (Sigma-Aldrich, Catalog # Z648043) with a MWCO of 100 kD.

### nrVL4619 scale-up production

Nucleofection, cell harvest, and lysis were performed as described for LY2 above except that the transfected plasmid encoded two copies of the nrVL4619 genome in tandem.

#### Iodixanol linear gradient and concentration

An iodixanol linear gradient was prepared by overlaying 19 mL of 20% iodixanol solution made in TN buffer with 19 mL of OptiPrep 60% iodixanol solution (Sigma-Aldrich) in 38.6 mL Ultra-Clear ultracentrifuge tubes (Beckman Coulter) and spinning on the Gradient Master (BioComp) at a 45-degree angle and a speed of 20 rpm for 16 minutes. Then the top 5 mL of iodixanol solution was replaced with 5 mL clarified lysate, and the tubes were ultracentrifuged at 32,000rpm and 20°C for 18 hours. After the O/N spin, 1-mL fractions were collected in 96 mL-deep well plates from the tops of the tubes. An aliquot of each fraction was used to measure refractive index using Refracto handheld refractometer (Mettler Toledo) as well as nrVL4619 titer, as per the protocol described above for the DNAse protected qPCR. Fractions of interest were determined based on the viral titer and density measurements. They were then pooled and concentrated ten-fold using the Amicon Ultra centrifugal filter units (Sigma-Aldrich, Catalog # Z648043) with a MWCO of 100 kD.

#### Size exclusion chromatography (SEC)

Prior to SEC, the sample was centrifuged at 12000 rpm for 1 minute. The supernatant was loaded onto a HiPrep 16/60 Sephacryl S-500 HR column (Cytiva) with buffer conditions at 50 mM Tris pH 8.0, 150 mM NaCl, and 0.01% poloxamer. The entire purification was performed at 4°C with a 1 mL/minute flow rate. The fractions with significant qPCR numbers were pooled and concentrated using Vivaspin 2, 10,000 MWCO PES concentrator (Sartorius, catalog # VS0201) and Nanosep centrifugal devices with Omega membrane at MWCO of 30K (Pall, catalog # OD030C34).

### Electron microscopy

To visualize virus particles, negative-stained TEM was conducted at Harvard Medical School using Jeol 1200 EX equipped with an AMT 2k CCD camera. 10 µl of sample was blotted on 400-mesh carbon support film (EMS CF400-Cu) for 30 seconds. After washing with double-distilled water for 30 seconds, the grid was stained by 1% of uranyl acetate for 10 seconds before imaging.

### *In vivo* studies

#### Care and use of animals

All mouse studies were approved and governed by the Laronde Institutional Animal Care and Use Committee. Female C57Bl/6J mice 8-12 weeks of age were obtained from Jackson Laboratories for use in these ocular studies.

#### Subretinal injections

Pupils were first dilated with one to two drops of 1% tropicamide/2.5% phenylephrine HCl (Tropi-Phen, Pine Pharmaceuticals). The mouse was subsequently anesthetized using an intraperitoneal injection of a ketamine/xylazine cocktail (100/10 mg/kg). One or two drops of 0.5% proparacaine (McKesson Corp.) were applied to the eye. An incision approximately 0.5 mm in length was made with a micro scalpel 1 mm posterior to the nasal limbus. A 33-g blunt-ended needle on a 5-μl Hamilton syringe was inserted through the scleral incision, posterior to the lens, toward the temporal retina until resistance was felt. 1 µl of either PBS, virus, or vector containing 0.1% of sodium fluorescein (AK-Fluor 10%, Akorn) was then injected slowly into the subretinal space. The eye was examined and the success of the subretinal injection was confirmed by visualizing the fluorescein-containing bleb through the dilated pupil with a Leica M620 TTS ophthalmic surgical microscope (Leica Microsystems, Inc). Eyes with significant hemorrhage or leakage of vector solution from the subretinal space into the vitreous were excluded from the study. After the procedure, 0.3% tobramycin ophthalmic ointment (Tobrex, Alcon) was applied to each treated eye and the mouse was allowed to recover from the anesthesia prior to being returned to its cage in the housing room.

#### Intravitreal injections

Pupils were first dilated with one to two drops of 1% tropicamide/2.5% phenylephrine HCl (Tropi-Phen, Pine Pharmaceuticals). The mouse was subsequently anesthetized using an intraperitoneal injection of a ketamine/xylazine cocktail (100/10 mg/kg). One or two drops of 0.5% proparacaine (McKesson Corp.) were applied to the eye. A 34-g beveled needle on a 5-μl Hamilton syringe was inserted 1 mm posterior to the nasal limbus, taking care not to damage the lens. 1 µl of either PBS, virus, or vector containing 0.1% of sodium fluorescein (AK-Fluor 10%, Akorn) was then injected slowly into the subretinal space. The eye was examined, and the success of the intravitreal injection was confirmed by visualizing the fluorescein-containing vitreous through the dilated pupil with a Leica M620 TTS ophthalmic surgical microscope (Leica Microsystems, Inc). Eyes with significant hemorrhage, lens damage, or leakage of vector solution outside of the eye were excluded from the study. After the procedure, 0.3% tobramycin ophthalmic ointment (Tobrex, Alcon) was applied to each treated eye and the mouse was allowed to recover from the anesthesia prior to being returned to its cage in the housing room.

#### Harvesting and processing of tissue samples for DNA extraction

Mouse eyes were dissected at indicated time points following subretinal or intravitreal injections (n = 5 for each time point). After enucleation, the retina and posterior eyecup (PEC) were separated and processed individually. These tissues were collected in tubes containing stainless steel beads and flash-frozen immediately. They were stored at -80°C until ready for homogenization. Frozen tissues were homogenized using Geno/Grinder 2010 (SPEX SamplePrep, LLC) at 1250 rpm for 30 seconds. Genomic DNA was isolated from homogenized tissues using the DNEasy Blood and Tissue Kit (Qiagen) according to the manufacturer’s instructions and quantified on Qubit Fluorometer using the Qubit DNA broad range Assay Kit (Thermo Fisher).

#### Quantitative PCR analysis

Genomic DNA was assayed by qPCR on the QuantStudio 5 – Real-Time PCR System (Thermo Fisher) using TaqMan Universal PCR Mastermix (Thermo Fisher). The sequence detection primers and the custom Taqman probe that were used in this study were synthesized by Integrated DNA Technologies (Table 1). All of the reactions including the DNA samples and different dilutions of a known quantity of the linearized mCherry or nrVL4619 plasmid standards were run in triplicate on the same plate. The standard curve method was used to calculate the amount of viral/vector DNA, which was normalized with the total amount of genomic DNA for each sample (quantified using Qubit as described above).

**TABLE 2.**
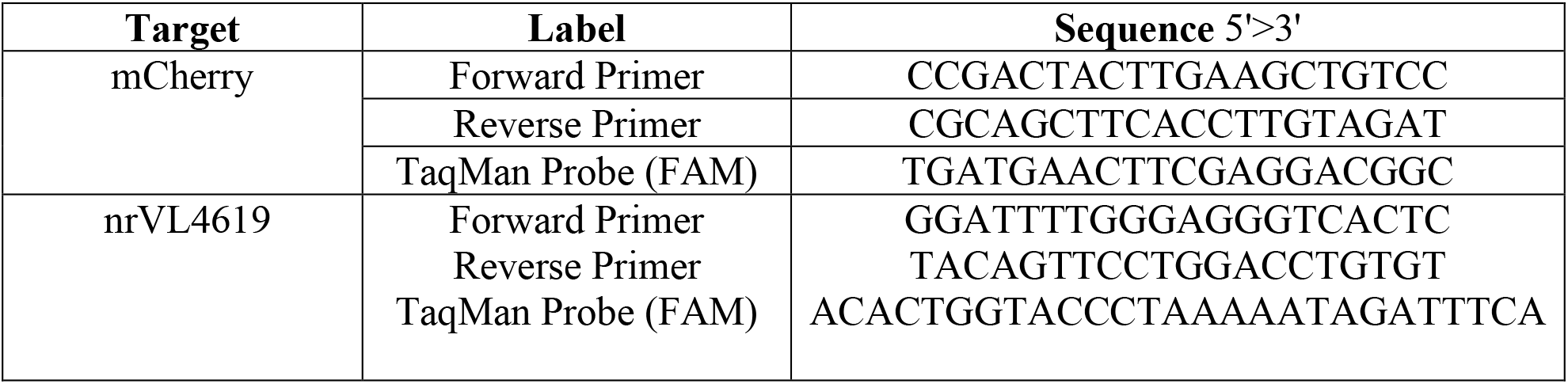
Primer and probes designed to quantify AAV2.mCherry and WT nrVL4619 in the DNA harvested from *in vivo* study tissue samples.

## RESULTS

### LY2 promoter is active in MOLT-4 cells

The viral load of anelloviruses in human plasma has been reported to be a hundred-fold lower than in whole blood, suggesting that cellular components of the blood harbor anelloviruses^38^, consistent with previous reports of lymphocytes being a major site of anellovirus replication^39–42^. Therefore, we examined whether anellovirus genes can be expressed in MOLT-4, a T-cell line derived from a patient with acute lymphoblastic leukemia^43^. For this, we synthesized a plasmid encoding two copies of the LY2 genome in a tandem arrangement. LY2 is a human anellovirus belonging to the *Betatorquevirus* genus that was previously sequenced from the pleural fluids of children hospitalized in France with parapneumonic empyema^37^. MOLT-4 cells electroporated with this plasmid were harvested at Days 1, 2, 3 and 4 post-transfection and analyzed for the detection of LY2 transcripts by RT-qPCR.

Previous anellovirus gene expression studies have described three major mRNA isoforms that are produced as a result of alternative splicing (Fig. 1A)^44^. For our analysis, we used a primer pair that would detect all 3 known isoforms of anellovirus transcripts. Expression of the GAPDH transcript was used for normalization. We were able to detect LY2 transcripts starting at Day 1 post-electroporation. Expression peaked at Day 3 post-transfection and was reduced at Day 4 (Fig. 1B). As expected, we did not detect any LY2 transcripts in untransfected MOLT-4 cells.

**FIGURE 1.**
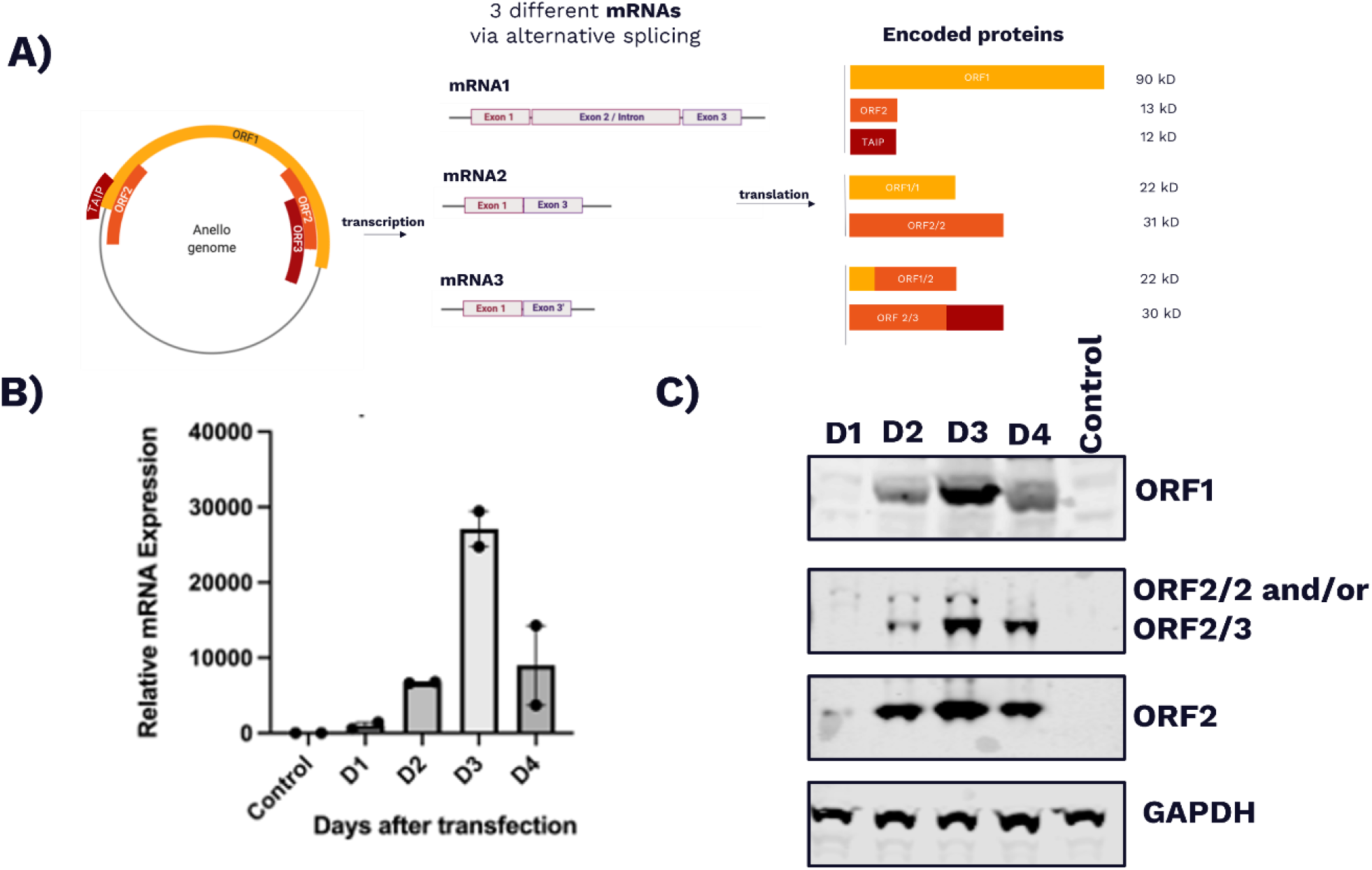
LY2 gene expression in MOLT-4 cells. A) Schematic of the single-stranded, circular DNA genome of an anellovirus, alternatively spliced to generate three different mRNAs encoding seven putative proteins of varying molecular weights as previously described. B) 10^7^ MOLT-4 cells were electroporated with 100 µg of a plasmid encoding two copies of the LY2 genome in tandem. RT-qPCR was performed at Days 1 (D1), 2 (D2), 3 (D3), and 4 (D4) post-electroporation to study the kinetics of the expression of LY2 transcripts over time. Untransfected MOLT-4 cells (control) were used as a negative control and GAPDH mRNA was used as a housekeeping gene for normalization. C) 10^7^ MOLT-4 cells were electroporated with 100 µg of a plasmid encoding two copies of the LY2 genome in tandem. Western blot analysis was performed at D1, D2, D3, and D4 post-electroporation to study the kinetics of the expression of LY2 proteins (ORF1, ORF2, ORF2/2 and/or ORF2/3) over time. GAPDH protein was used as a loading control.

Having detected LY2 transcripts, next we performed a similar time-course experiment to determine LY2 protein expression. Antibodies to detect the putative capsid protein, ORF1, as well as ORF2 and its variants were generated. As shown in Figure 1C, ORF1 is detectable beginning on Day 2 post-transfection, peaks on Day 3, and is reduced beyond Day 3. As the epitope used to generate the anti-ORF1 antibody is specific to the jelly roll domain of ORF1, it cannot detect isoforms such as ORF1/1 and ORF1/2. The anti-ORF2 antibody that we generated can detect all three isoforms of ORF2, including ORF2, ORF2/2, and ORF2/3. The predicted molecular weights for ORF2, ORF2/2, and ORF2/3 are 17, 31, and 30 kDa, respectively. Since ORF2/2 and ORF2/3 are nearly equal in molecular weight, it is challenging to distinguish them based on migration on SDS-PAGE gel in a denatured state. Like ORF1, the expression of ORF2 and its isoforms also peaked on Day 3 post-transfection and was reduced thereafter (Fig. 1C). Collectively, these results suggest that the LY2 promoter is active in MOLT-4 cells, enabling transcription and translation of anellovirus genes in this human cell line.

### MOLT-4 is permissive for the replication of the LY2 anellovirus

Having detected LY2 gene expression in MOLT-4 cells, next we tested whether the cell line is permissive for replication of the anellovirus genome. We nucleofected plasmids to encode either a single copy of the LY2 genome or two copies of the LY2 genome in tandem. The cells were harvested four days post-nucleofection, followed by DNA extraction. Extracted DNA was either left untreated or treated with a restriction enzyme that digests the plasmid backbone once, a restriction enzyme that digests the LY2 genome once, or DpnI. DNA replicated in bacterial cells contains methylated adenine and therefore is sensitive to digestion with DpnI. On the other hand, DNA that replicates in eukaryotic cells lacks methylated adenine and therefore is resistant to digestion with DpnI. Hence, DpnI digestion can be used to distinguish between the transfected genome and any genome that may have replicated in MOLT-4 cells. Untreated and treated DNA samples were subjected to Southern blot analysis using probes designed to specifically detect the LY2 genome.

For DNA extracted from the sample transfected with a tandem LY2 genome-containing plasmid (Fig. 2, Sample #5), we detected a band of the same size as would be expected for a unit-length, double-stranded LY2 genome. This band was insensitive to digestion with an enzyme that cuts the plasmid backbone but became linear when treated with an enzyme that cuts the LY2 genome once. These observations suggest that this band represents the LY2 genome and not the plasmid backbone. Furthermore, this band was resistant to treatment with DpnI, indicating that the LY2 genome replicated in the transfected MOLT-4 cells. Overall, based on the patterns of banding observed for samples and controls, we conclude that MOLT-4 cells transfected with a tandem LY2 genome-containing plasmid are permissive for replication of the anellovirus and lead to a unit-length, replicated anellovirus genome.

**FIGURE 2.**
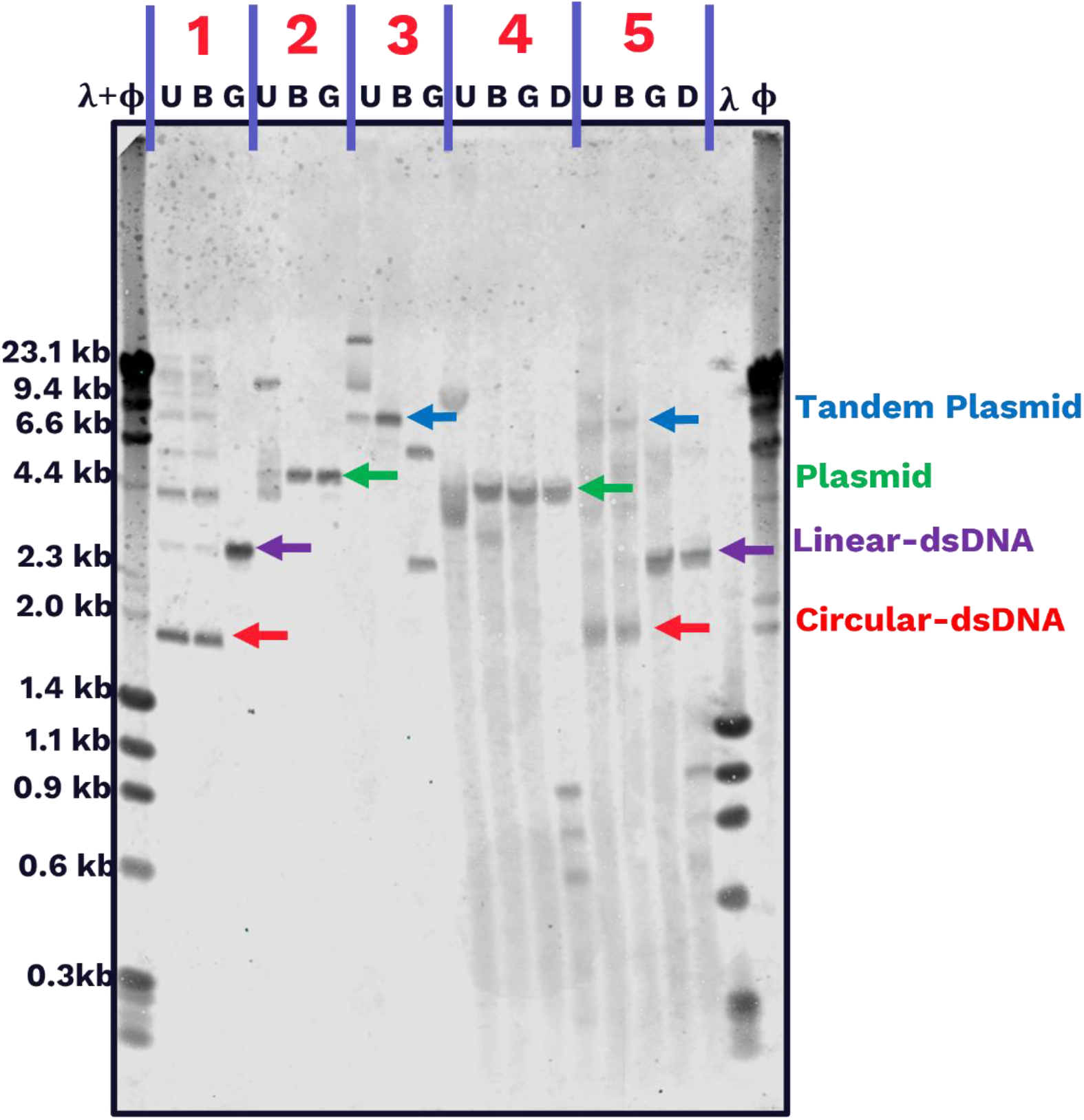
MOLT-4 cells are permissive for LY2 replication. 10^7^ MOLT-4 cells were transfected with 25 µg of either a plasmid encoding a single copy of the LY2 genome (Sample #4) or a plasmid encoding two copies of the LY2 genome in tandem (Sample #5). Total genomic DNA was harvested from the cells at four days post-transfection and was either untreated (U) or treated with an enzyme intended to digest the plasmid backbone once (B), an enzyme intended to digest the LY2 genome once (G), or DpnI restriction enzyme (D). As controls, enzyme treatments were also performed in parallel on an in-vitro-circularized LY2 genome (Sample #1), a plasmid containing a single copy of the LY2 genome (Sample #2), and a plasmid containing two copies of the LY2 genome in tandem (Sample #3). Southern blot analysis was performed on the digested samples using probes specific against the LY2 genome. The expected sizes for the plasmid containing tandem LY2 genomes, the plasmid containing a single LY2 genome, a unit-length double-stranded linear genome, and a unit-length double-stranded circular genome are indicated by blue, green, purple, and red arrows, respectively.

A DpnI-resistant band was also detected for the sample transfected with a single LY2 genome-containing plasmid (Fig. 2, Sample #4). When this sample was treated with a restriction enzyme that digests either the plasmid backbone once or the LY2 genome once, we detected a band of the same size as would be expected for a linear plasmid containing the entire LY2 genome. This result suggests that the single LY2 genome-containing plasmid can also replicate in MOLT-4 cells but does not yield a detectable unit-length anellovirus genome. To our knowledge, this is the first study to conclusively demonstrate the replication of a unit-length, circular, human anellovirus genome in a human cell line using recombinant DNA as an input material.

### MOLT-4 cells are permissive for LY2 packaging

Since we demonstrated that MOLT-4 cells are permissive for LY2 replication, we next tested whether LY2 particles can be produced in this human cell line. We nucleofected MOLT-4 cells with either a plasmid containing a qPCR amplicon of LY2 (LY2 non-rep) or an *in vitro* circularized, double-stranded LY2 genome (LY2 IVC). Four days after nucleofection, cells were harvested, lysed by two rounds of freeze-thawing, treated with Benzonase, clarified, and subjected to isopycnic ultracentrifugation using CsCl linear gradient. 1-mL fractions of CsCl linear gradient were collected from the bottom of the tube. Each fraction was analyzed for its density as well as for titers of LY2 titer using a DNAse-protected qPCR assay.

As shown in Fig. 3, the sample transfected with LY2 IVC showed a peak viral titer in fractions with a density of approximately 1.32 g/cm^3^. This density is consistent with the previously described density of anelloviruses in CsCl^45,46^. In contrast, the sample transfected with negative control had no detectable viral titer in any fractions. These results conclusively demonstrate that the MOLT-4 cell line is permissive not only for LY2 gene expression and replication, but also for production of LY2 particles. While isopycnic ultracentrifugation to analyze the density of anellovirus particles in CsCl has been previously reported for wild-type anellovirus particles isolated from human specimens, our study is the first to obtain such results for anellovirus particles produced *in vitro* using a recombinant viral genome.

**FIGURE 3.**
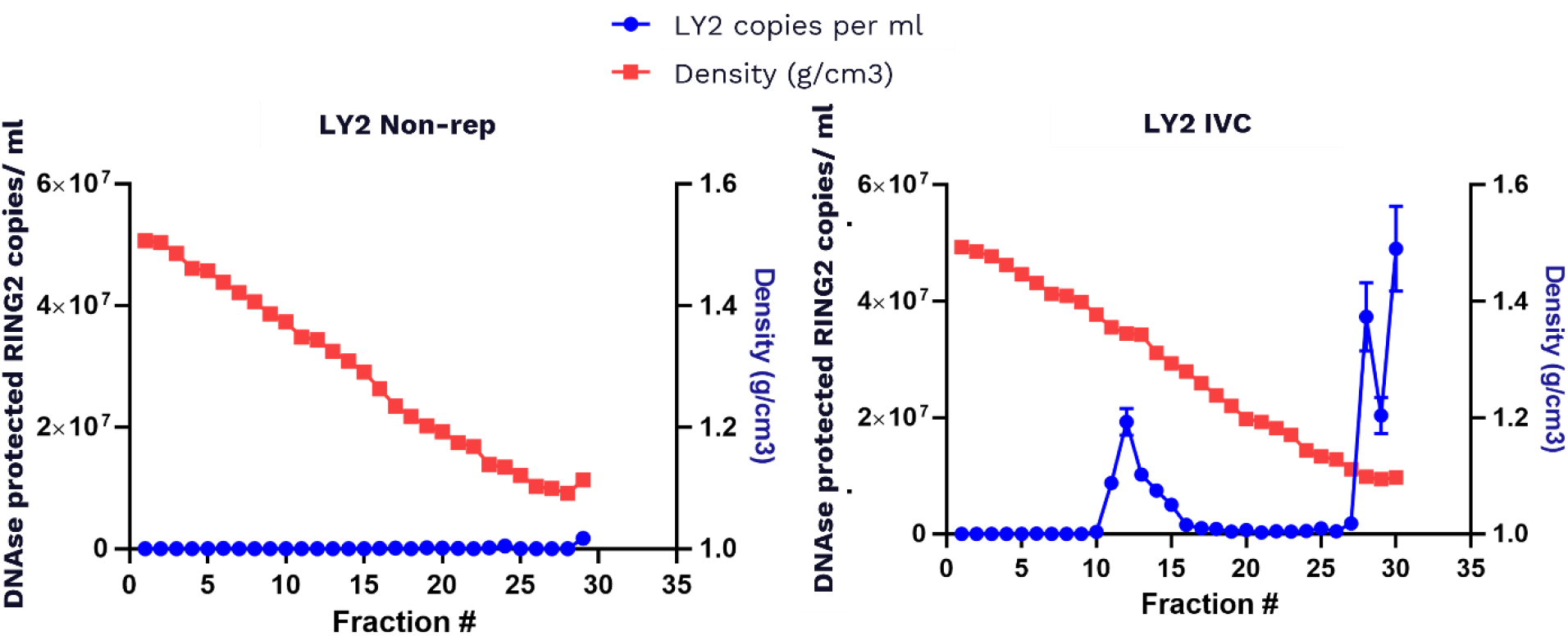
Packaging of LY2 particles in MOLT-4 cells. 10^7^ MOLT-4 cells were transfected with 25 µg of either a negative control plasmid (non-rep) or an *in vitro* circularized genome of LY2 (IVC). Cells were harvested 4 days post-transfection, lysed, and treated with Benzonase. Clarified lysate was then subjected to isopycnic centrifugation using CsCl linear gradient. Each collected fraction was analyzed for density (plotted in red) and viral titer (plotted in blue). A representative linear gradient profile is shown.

### Production of LY2 in MOLT-4 cells is dependent on viral protein expression

To assess whether the production of LY2 particles is dependent on viral protein expression, we created LY2 mutant genomes in which either all 3 ORF1 variants including ORF1, ORF1/1, and ORF1/2 were knocked out (ORF1 KO), or all 3 ORF2 variants including ORF2, ORF2/2, and ORF2/3 were knocked out (ORF2 KO). These mutant genomes were generated by inserting premature stop codons into the open reading frames, as described in materials and methods.

To confirm successful knockout of the target proteins, plasmids encoding a single copy of either the wild-type LY2 genome, the ORF1 KO genome, or the ORF2 KO genome were transfected into MOLT-4 cells. Western blot analysis was performed at 2 days and 3 days post-transfection. As expected, the ORF1 KO mutant did not express any detectable ORF1 protein (Fig. 4A). Similarly, the ORF2 KO mutant did not express any detectable levels of ORF2 protein or its isoforms (Fig. 4B). To test whether these mutants can produce LY2 virus particles, MOLT-4 cells were transfected with either a plasmid encoding two copies of the wild-type LY2 genome in tandem (WT LY2 tandem), an *in vitro* circularized ORF1 knockout genome (ORF1 KO IVC), or an *in vitro* circularized ORF2 knockout genome (ORF2 KO IVC) or were co-transfected with ORF1 KO IVC and ORF2 KO IVC. Samples were assayed for LY2 production using isopycnic CsCl step gradient. As expected, WT LY2 tandem produced LY2 particles (Fig. 4C). Knocking out the expression of ORF1 and its variants or ORF2 and its variants significantly disrupted the ability to produce the virus in MOLT-4 cells. Interestingly, the mutant genomes were able to trans-complement each other, which was expected assuming co-transfection of the same cells, given that ORF1 KO IVC can produce ORF2 and its variants while ORF2 KO IVC can produce ORF1 and its variants. Overall, these findings demonstrate that the production of LY2 particles in transfected MOLT-4 cells is dependent on viral protein expression.

**FIGURE 4.**
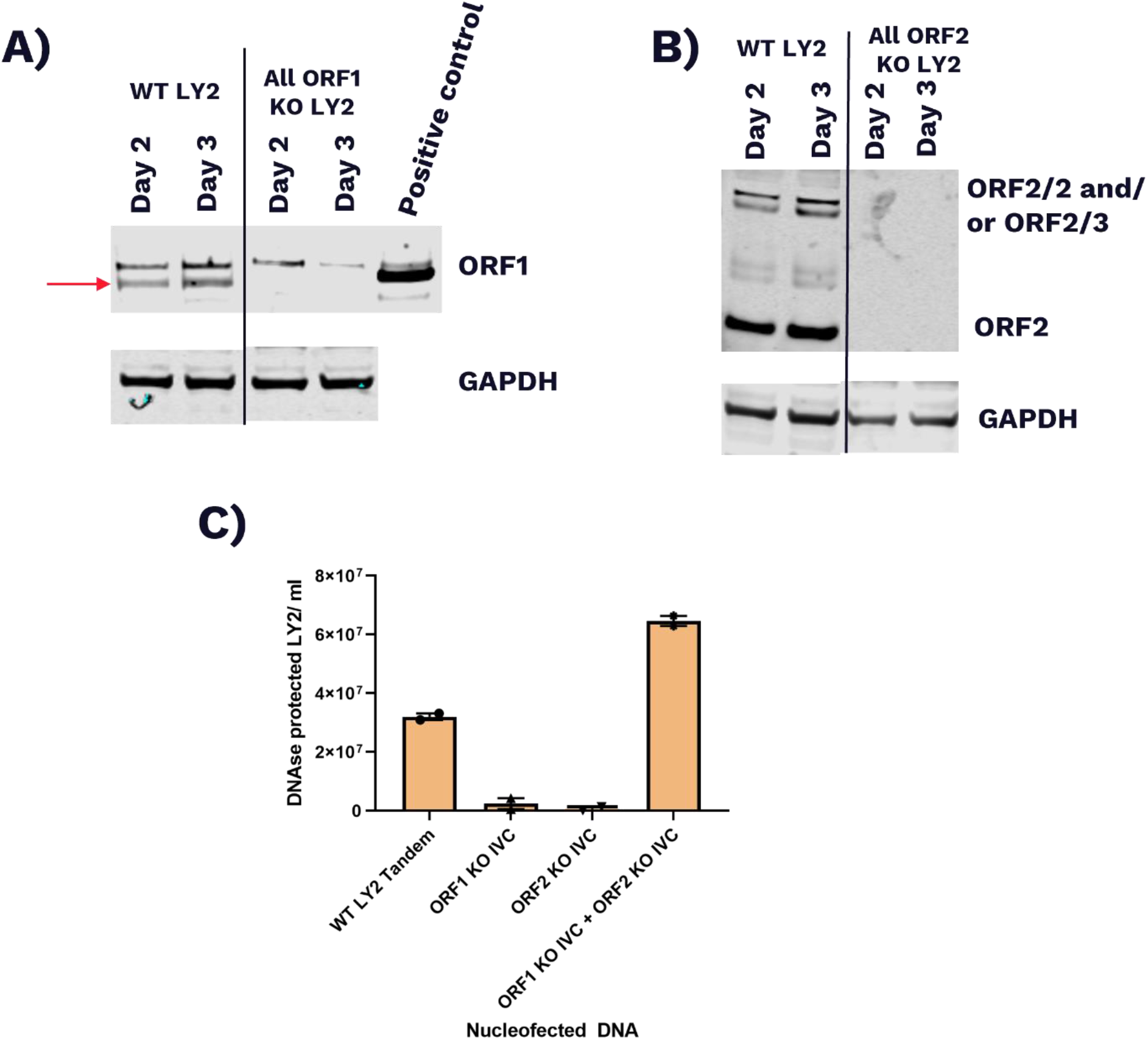
Packaging of LY2 particles in MOLT-4 cells is ORF1- and ORF2-dependent. 10^7^ MOLT-4 cells were electroporated with 100 µg of plasmid encoding either the wild-type LY2 genome (WT), a LY2 genome in which the expression of all ORF1 variants has been knocked out (All ORF1 KO), or a LY2 genome in which the expression of all ORF2 variants has been knocked out (All ORF2 KO). Cells were harvested either 2 or 3 days post-transfection. A) Western blotting was performed to detect ORF1 protein. Purified ORF1 was used as a positive control. The ORF1 band is shown using a red arrow. GAPDH was used as a loading control. The black line denotes where the blot was cut out to remove unnecessary lanes. B) Western blotting was performed to detect ORF2 and its variants. GAPDH was used as a loading control. The black line denotes where the blot was cut out to remove unnecessary lanes. C) 10^7^ MOLT-4 cells were transfected with either 25 µg of a plasmid encoding two copies of the LY2 genome in tandem (WT LY2 tandem), 25 µg of an *in vitro* circularized genome of LY2 in which the expression of all ORF1 variants has been knocked out (ORF1 KO IVC), or 25 µg of an *in vitro* circularized genome of LY2 in which the expression of all ORF2 variants has been knocked out (ORF2 KO IVC), or were co-transfected with 12.5 µg each of ORF1 KO IVC and ORF2 KO IVC. Cells were harvested 4 days post-transfection, lysed, and treated with Benzonase. Clarified lysates were then subjected to isopycnic centrifugation using CsCl step gradient to isolate proteins within a range of density from 1.2 to 1.4 g/cm^3^. The isolated fraction was dialyzed to remove CsCl and then was used to perform DNAse-protected qPCR

### Transmission electron microscopy (TEM) analysis of LY2 anellovirus

Visualization of recombinant anellovirus particles has been previously performed only for chicken anemia virus, an avian virus in the *Anelloviridae* family^47^. To further characterize and advance our understanding of the structural biology of human anelloviruses, we analyzed LY2 by TEM. A schematic of the production and purification methodology of LY2 is depicted in Fig. 5A. Briefly, MOLT-4 cells were transfected with a plasmid containing two copies of the LY2 genome in tandem and harvested four days post-transfection. Harvested cells were lysed and treated with Benzonase and detergent. Cell lysates were clarified to remove any cellular debris and subjected to CsCl step gradient to concentrate the virus particles followed by overnight dialysis to remove any CsCl. Dialyzed material was subjected to CsCl linear gradient followed by fractionation. Each fraction was analyzed for density and viral titer.

**FIGURE 5.**
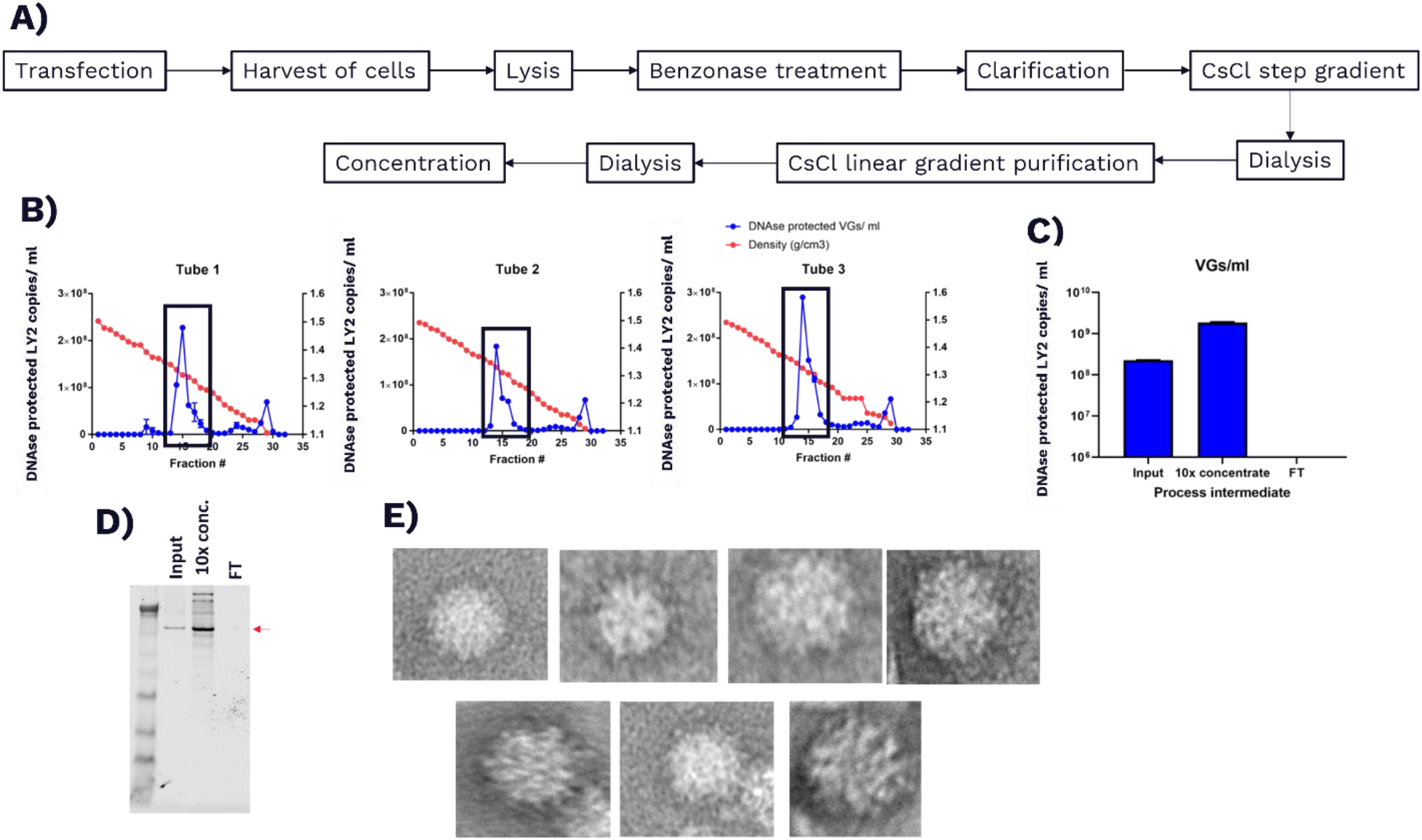
Electron microscopy detection of LY2 particles produced in MOLT-4 cells. A) Schematic of the production and purification of LY2 particles from MOLT-4 cells. B) 2x 10^9^ MOLT-4 cells were transfected with 2 mg of a plasmid containing two copies of the LY2 genome in tandem. Cells were harvested 4 days post-transfection, lysed, treated with Benzonase, clarified, concentrated using CsCl step gradient, and dialyzed. Dialyzed material was subjected to isopycnic centrifugation using CsCl linear gradient. Each fraction of linear gradient was analyzed for density and viral titer. Three representative linear gradient profiles are shown. Fractions with the expected density of LY2 (shown with black box) were pooled together and concentrated 10× using Centricon centrifugal filter units. C) The viral titers in the pooled material (input), concentrated material, and flow-through (FT). D) Western blot analysis to detect capsid protein ORF1 in the pooled material (input), concentrated material, and FT. E) Representative TEM images of concentrated LY2 particles.

As expected, all linear gradient tubes had a peak for DNAse-protected LY2 titer at the expected density of 1.32 g/cm^3^ ^45^. Representative profiles for density against viral titer in each fraction of the linear gradient are shown in Fig. 5B. Next, we pooled the fractions in the peak for all 12 tubes of linear gradient, dialyzed it overnight to remove any CsCl, and concentrated the volume tenfold using diafiltration. As we used 100 kD cutoff diafiltration units, we were able to concentrate the titer of LY2 particles (Fig. 5C). Concomitantly, an increase in the titer of capsid protein ORF1 as assessed by Western blot was observed as expected (Fig. 5D). Negative-staining TEM of this purified virus preparation revealed multiple LY2 particles (Fig. 5E). The observed particles we detected were consistent with the expected ∼30nm-diameter viral particles.

### Discovery of an anellovirus in human retinal pigment epithelial (RPE) cells

Anelloviruses have previously been isolated from numerous human non-blood tissues, such as the bone marrow, liver, and conjunctival surface and vitreous fluid of the eye^48–51^. We investigated the specific anellovirus lineages present in four ocular tissues (cornea, macula, sclera and retina pigment epithelia) from the same subject using our AnelloScope platform^14^. We recovered several anellovirus genomes, across all three genera, from half of the investigated ocular tissues and successfully isolated a putative full-length circularized genome designated as nrVL4619 (Fig. 6A).

**FIGURE 6:**
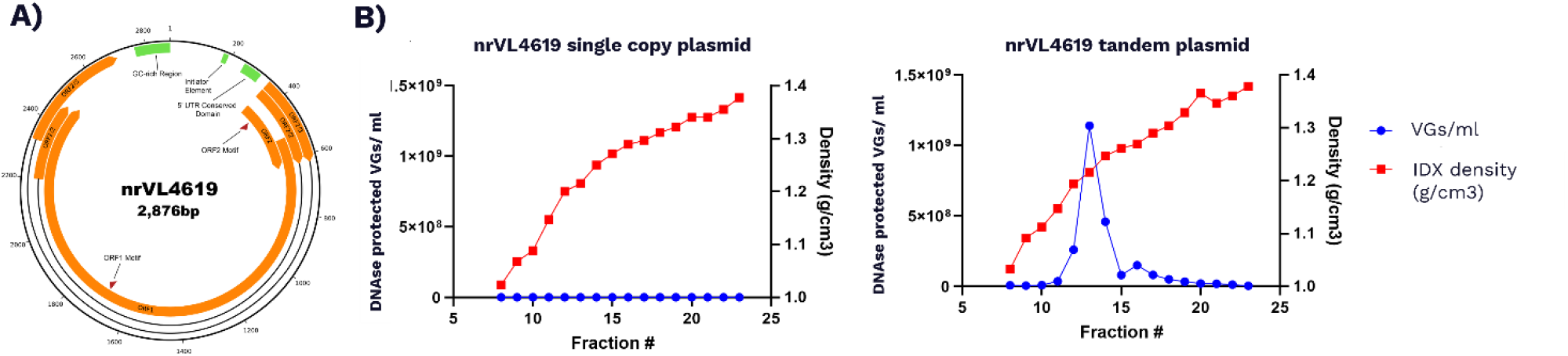
Recovery of a circularized, complete anellovirus genome from RPE cells and its rescue in MOLT-4 cells. A) Schematic of the fully annotated, circularized genome of nrVL4619 recovered from a dissected RPE tissue. ORF1 and ORF2 were computationally annotated, while ORF2/2 and ORF2/3 were manually curated. B) 10^7^ MOLT-4 cells were transfected with 50 µg of either a plasmid encoding a single copy of the nrVL4619 genome or a plasmid encoding two copies of NRVL4619 in tandem. Cells were harvested 4 days post-transfection, lysed, and treated with Benzonase. Clarified lysate was then subjected to isopycnic centrifugation using iodixanol linear gradient. Each collected fraction was analyzed for density (plotted in red as g/cm^3^) and viral titer (plotted in blue as viral genomes (vgs)/ml). A representative linear gradient profile is shown.

To explore the diversity of anelloviruses in the four eye tissues (cornea, macula, sclera and retina pigment epithelia), we conducted two deep short-read sequencing runs (with and without bead-baited target enrichment) and one shallow long-read sequencing run to recover an appropriate amount of genomic anellovirus data. An aggregate of 28.71 Gbp of short-read sequence data and 1.41 Gbp of long-read sequence data were generated across all sequencing runs, of which 3.71 Gbp and 269.3 Mbp of sequence data were classified as anellovirus from short- and long-read sequencing runs, respectively. Strikingly, when examining the two short-read sequencing runs, the run utilizing our bead-baited target enrichment protocol contributed 99.9% of those reads identified as anellovirus. These findings highlight the difficulty in isolating anellovirus genome data from non-blood tissue samples and the need for targeted approaches that both amplify the amount of anellovirus in samples and reduce the amount of host background being sequenced.

We next attempted to recover complete, circularized, high-quality anellovirus genomes from the RPE genomic data generated. We searched through the existing short-read-assembled contigs to find suitable candidates that were near genome length and contained the expected ORF1, ORF2, ORF3, 5’ UTR, and 3’ GC-rich region features, producing a *Betatorquevirus* candidate designated nrVL4619*-short*. To confirm the completeness of nrVL4619-*short*, we examined the paired long-read data for any sequences with at least 90% sequence similarity. We found 7 long reads, that were further assembled and error-corrected to produce a contig designated nrVL4619-*long*. When comparing nrVL4619-*short* to nrVL4619-*long*, we observed 98.4% similarity, translating to a decrease of 4% in error rate from the pre-corrected long-read sequences. To further improve accuracy, we leveraged the accompanying short-read genomic data by mapping these reads (485,785 mapped reads, 4,275 average coverage) to nrVL4619-*long* over three rounds of additional error-correction, resulting in a final sequence at 99.9% similarity when compared to nrVL4619-*short*.

To resolve the 0.1% divergence observed between nrVL4619-*long* and nrVL4619-*shor*t, found at four positions, we employed high-quality Sanger sequencing. We recovered two overlapping Sanger reads (both forward and reverse) for each of the four sites identified and resolved inconsistencies between nrVL4619-*short* and -*long* at three of the four sites. At the fourth site, located in the ORF1 capsid protein at positions 1330 and 1340, we found no clear consensus sequence established from the Sanger data. Examining all the data generated at this fourth site indicated the existence of an RPE-specific nrVL4619 consensus sequence. In producing a final high-quality sequence of nrVL4619, we pursued the RPE dominant consensus sequence.

### Production of nrVL4619 virus particles *in vitro*

Whereas LY2 had been previously sequenced from human pleural effusion samples and reported in the literature, nrVL4619 is an anellovirus that we isolated from human retinal pigmental epithelium as described above. To test whether nrVL4619 can be produced *in vitro* like LY2, we electroporated MOLT-4 cells with either a plasmid encoding a single copy of the nrVL4619 genome or a plasmid encoding two copies of the nrVL4619 genome in tandem. Four days after electroporation, cells were harvested, lysed by two rounds of freeze-thawing, treated with Benzonase, clarified, and subjected to isopycnic ultracentrifugation using iodixanol linear gradient. 1-mL fractions of linear gradient were collected from the top of the tube. Each fraction was analyzed for its density as well as for titers of nrVL4619 using a DNAse-protected qPCR assay. As shown in Fig. 6B, the sample transfected with nrVL4619 tandem plasmid showed an enrichment of viral genomes in fractions with a density of approximately 1.21 g/cm^3^, suggestive of successful production of nrVL4619 in MOLT-4 cells. On the other hand, we detected significantly lower to no titers of nrVL4619 in any of the linear gradient fractions tested in the sample transfected with a single copy of the nrVL4619 genome.

To test whether we can visualize nrVL4619 virus particles, we scaled up production and purification as summarized in Fig. 7A. Briefly, the processed lysates of the transfected cells were subjected to a two-step purification including isopycnic centrifugation using an iodixanol linear gradient followed by size exclusion chromatography (SEC). A clear peak for the nrVL4619 titer was detected in SEC fractions in which anellovirus particles would be expected to migrate (Fig. 7B). When these fractions were pooled and concentrated for TEM analysis, we detected multiple nrVL4619 particles as shown in Fig. 7C and 7D. The morphology of these nrVL4619 particles was consistent with the morphology of LY2 particles (Fig. 6E).

**FIGURE 7.**
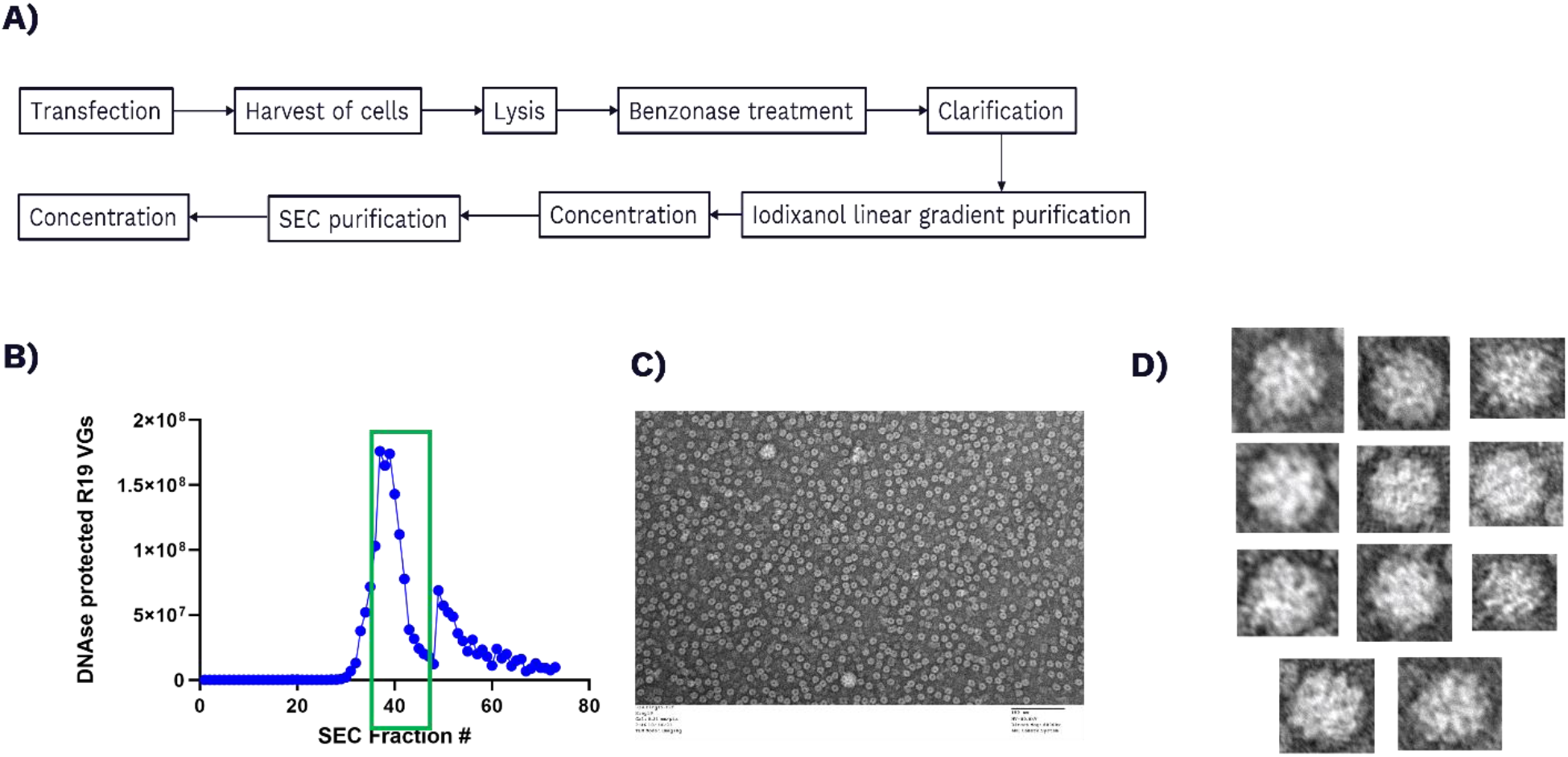
Electron microscopy detection of nrVL4619 particles produced in MOLT-4 cells. A) Schematic of the production and purification of nrVL4619 particles from MOLT-4 cells. B) DNAse-protected qPCR assay of fractions from SEC of purified nrVL4619. C and D) Representative TEM images of concentrated NRVL4619 particles.

### nrVL4619 particles demonstrate infectivity *in vivo*

Since nrVL4619 genome was isolated from human retinal epithelium, we hypothesized that it would have tropism for eye tissue. To examine this, we tested its infectivity and tropism in the eye *in vivo*. Mice were injected either subretinally or intravitreally with PBS, purified WT nrVL4619, or dose-matched AAV2.mCherry as shown (Fig. 8A). Eyes were harvested and separated into the neuroretina (which contains the photoreceptor, bipolar, and ganglion cells) and the posterior eye cup (PEC, which contains the retinal pigmented epithelium, choroid, and sclera). DNA was harvested from these tissues followed by qPCR analysis to detect nrVL4619 and AAV2.mCherry genomes. nrVL4619 demonstrated infectivity in both the neuroretina and PEC on Days 7 and 21 following either subretinal or intravitreal injections (Fig. 8B). nrVL4619 demonstrated superior targeting of the neuroretina and PEC following subretinal injection compared to AAV2.mCherry. This increase in infectivity was observed on both Days 7 and 21. For intravitreal injections, nrVL4619 also demonstrated superior infectivity in the PEC on Days 7 and 21 compared to AAV2.mCherry. Additionally, we detected nrVL4619 infectivity in the neuroretina following intravitreal delivery on Days 7 and 21. These data suggest that nrVL4619, which was isolated from the RPE of a human donor, demonstrates superior targeting of the PEC than AAV2.

**FIGURE 8.**
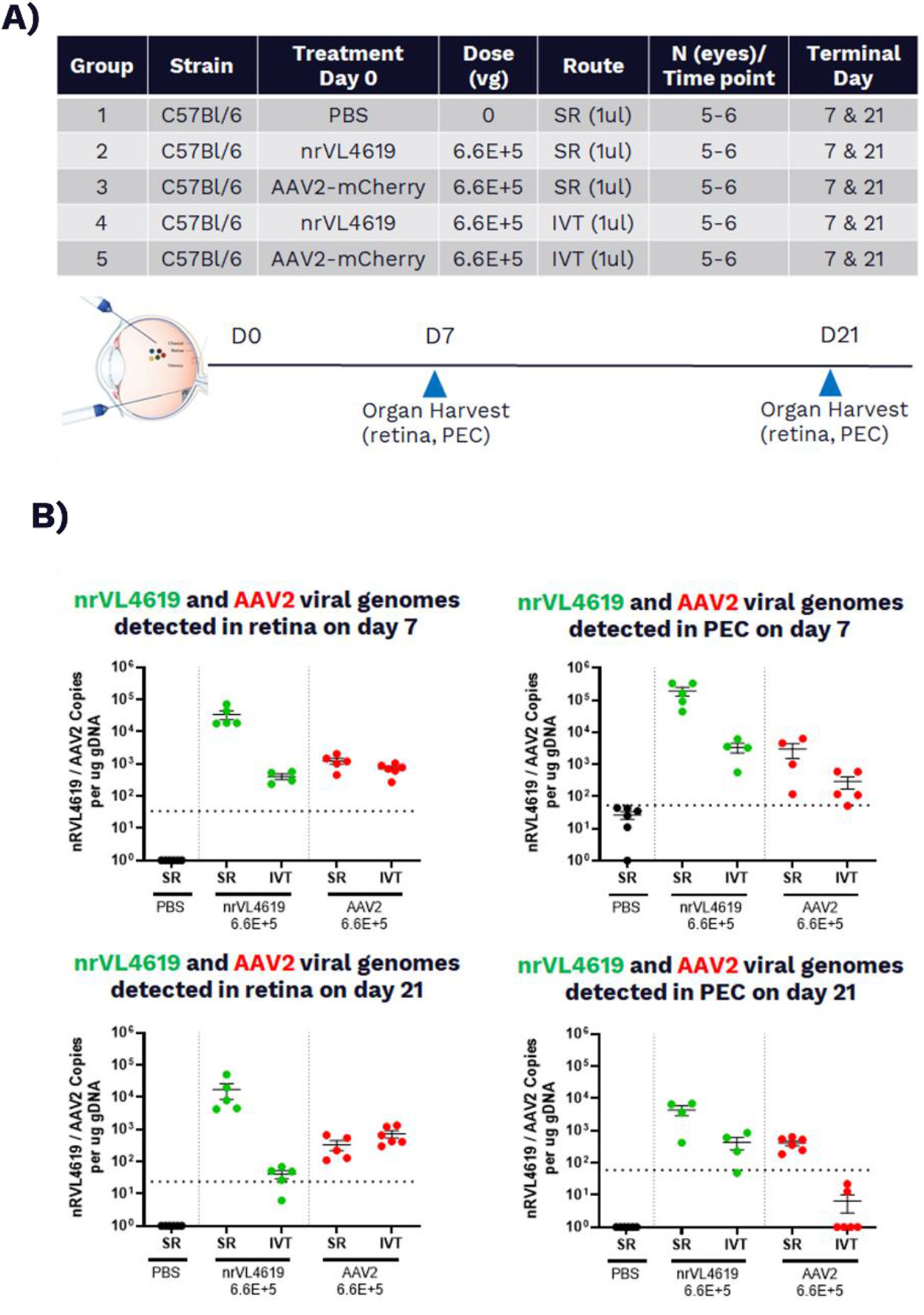
nrVL4619 infectivity in mouse retina and posterior eye cup (PEC). A) Table describing various strain of mouse, treatments, virus/vector doses, routes of administration, numbers of animals per group, and time points for the *in vivo* study. Bottom panel shows a schematic of the anatomy of a mouse eye as well as study design. (B) Vector/virus genome copies present in the neuroretina or PEC, as assessed by qPCR in the harvested DNA from mouse eyes injected intravitreally (IVT) or subretinally (SR) once with either PBS, 6.6E+5 vg of Ring 19, or dose-matched AAV2.mCherry. N = 5-6 eyes/group.

## DISCUSSION

Anelloviruses are a large and diverse family of viruses comprising the majority of the human virome. Anellovirus infections are acquired during infancy and detected throughout the lives of healthy individuals, suggesting they have evolved to avoid clearance by and live in harmony with the human immune system^10^. Previous independent studies as well as our own work have demonstrated the recovery of anelloviral sequences from many types of human tissues and biological samples^3–6^. In addition to evading the host immune system, anelloviruses might be leveraging their diversity to achieve broad tropism for a range of tissue and cell types. The non-pathogenic and weakly immunogenic nature of anelloviruses may overcome a major limitation of current vectors by enabling repeat dosing of gene therapies, while specific tropism of the diverse lineages of anelloviruses may enable development of an anellovirus-based gene therapy delivery platform capable of addressing diseases across multiple therapeutic areas.

Understanding the basic properties of these viruses further will offer new insights into their biology and unlock our ability to harness their biology for therapeutic applications. Previous studies generally have been limited to the detection and sequencing of anellovirus nucleic acids in blood or other specimens. We recently have reported on comprehensive efforts to use genomics, computational biology, and a series of biochemical and biophysical techniques to elucidate the diversity, transmission, structure, and immunogenicity of the anellovirus family. For example, we developed a proprietary AnelloScope technology to specifically enrich and recover sequences of anelloviruses from human tissue. We used this technology to demonstrate in a blood-transfusion cohort that anelloviruses can be transmitted from donors and can persist for at least nine months in recipients^14^. Recently, we used a phage immunoprecipitation sequencing (PhIP-Seq) assay to comprehensively profile antibody responses to the human anellome^52^. In this study, we introduce an *in vitro* system for the production of human anelloviruses, which will enable further basic research into anelloviruses, the knowledge from which can be used to develop anellovirus-based gene therapies.

As anelloviruses have been previously reported to reside and replicate in immune cells within the human body, we selected a T-cell-derived cell line of human origin, MOLT-4, and tested its permissiveness for the production of a *Betatorquevirus* LY2 originally described by a group in France. LY2 gene expression in MOLT-4 cells was observed, with anellovirus mRNA and protein levels peaking on the third day after transfection (Fig. 1). When MOLT-4 cells were transfected with a plasmid encoding tandem copies of the LY2 genome, replication of a unit-length, circular, double-stranded form of the viral genome was evident (Fig. 2). Notably, a plasmid encoding a single copy of the LY2 genome also can replicate in MOLT-4 cells but does not lead to virus rescue, presumably due to lack of the formation of unit-length circular genome (Fig. 2). The design of the tandem plasmid and its repetitive sequences may lead to genetic recombination and the generation of some species of the genome that are unit-length, circular, and double-stranded, which are then replicated by the virus and host replication machinery. Studies to determine the function of each of the anellovirus-encoded proteins as well as which host factors are involved in the viral life cycle represent an avenue for additional area of research that we are currently actively pursuing.

MOLT-4 cells transfected with the tandem plasmid or an *in vitro* circularized, double-stranded form of the LY2 genome were also permissive for the production of LY2 virus particles, as shown by peak viral titers in CsCl-based density gradient fractions (Fig. 3, Fig. 5) and reinforced by TEM analysis showing multiple particles with an icosahedral capsid (Figure 5). Our study is the first to document the production of a human anellovirus using recombinant viral DNA genome.

Viral production in MOLT-4 cells was not observed when either ORF1 and its isoforms or ORF2 and its isoforms were knocked out via site-directed mutagenesis of the LY2 genome, implying that expression of the combination of these proteins is critical for virus production. This conclusion is supported by the observation that viral production was restored when the cells were co-transfected with the ORF1 knockout and ORF2 knockout mutant genomes (Figure 4), indicating successful trans-complementation between the ORF2 proteins expressed by the ORF1 knockout and the ORF1 proteins expressed by the ORF2 knockout.

To assess whether other anelloviruses besides LY2 can be produced in MOLT-4 cells, we transfected MOLT-4 cells with a plasmid encoding tandem copies of nrVL4619. nrVL4619 is a separate *Betatorquevirus* that we isolated from the human retinal pigmental epithelium. Similar to LY2, we were able to detect production of nrVL4619 as assessed by isopycnic centrifugation and TEM (Fig. 6). When nrVL4619 was intravitreally or subretinally injected into mice, the anellovirus was shown to infect the retina as well as the posterior eye cup. These findings demonstrate tropism of nrVL4619 for ocular tissue and provide proof-of-concept for the use of the AnelloScope technology to discover anelloviruses with tropism for a tissue of interest.

In summary, we have established an *in vitro* cell-based recombinant system that is capable of producing human anelloviruses, a family of commensal viruses that are detectable in a variety of human tissues with high prevalence but have been understudied in the past. Using the MOLT-4 cell line to robustly express natural or engineered anelloviruses enables comprehensive evaluation of their biophysical and biochemical properties, their infectivity *in vitro* and *in vivo*, and their immune response profile. This study also represents an important step toward the application of non-pathogenic, weakly immunogenic anelloviruses as vectors for re-dosable gene therapies targeting a wide array of diseases.

## ACKNOWLEDGEMENTS

This study utilized TTVS research materials obtained from the NHLBI Biological Specimen and Data Repository Information Coordinating Center and does not necessarily reflect the opinions or views of the TTVSor the NHLBI.

## COMPETING INTEREST STATEMENT

DMN, MT, GB, CP, EO, JY, CAA, AB, DV, PT, JC, SL, KS, HS, AM, YC, TO, NLY, RJH, and SD are employees of and hold equity interests in Ring Therapeutics. TO and RJH are affiliated with Flagship Pioneering, which also holds equity interests in Ring Therapeutics. KL, CS, NA, and FD also hold equity interests in Ring Therapeutics.

## FUNDING

This study was funded by Ring Therapeutics.

## REFERENCES

1. Takahashi, K. Partial ∼2.4-kb sequences of TT virus (TTV) genome from eight Japanese isolates: diagnostic and phylogenetic implications. Hepatol Res 12, 111–120 (1998).

2. Nishizawa, T. et al. A novel DNA virus (TTV) associated with elevated transaminase levels in posttransfusion hepatitis of unknown etiology. Biochem Bioph Res Co 241, 92–97 (1997).

3. Takahashi, K., Iwasa, Y., Hijikata, M. & Mishiro, S. Identification of a new human DNA virus (TTV-like mini virus, TLMV) intermediately related to TT virus and chicken anemia virus. Arch Virol 145, 979–993 (2000).

4. Kaczorowska, J. & Hoek, L. van der. Human Anelloviruses: diverse, omnipresent and commensal members of the virome. Fems Microbiol Rev 44, 305–313 (2020).

5. Hijikata, M., Takahashi, K. & Mishiro, S. Complete Circular DNA Genome of a TT Virus Variant (Isolate Name SANBAN) and 44 Partial ORF2 Sequences Implicating a Great Degree of Diversity beyond Genotypes. Virology 260, 17–22 (1999).

6. Okamoto, H. et al. Species-specific TT viruses in humans and nonhuman primates and their phylogenetic relatedness. Virology 277, 368–378 (2000).

7. Ninomiya, M., Takahashi, M., Nishizawa, T., Shimosegawa, T. & Okamoto, H. Development of PCR Assays with Nested Primers Specific for Differential Detection of Three Human Anelloviruses and Early Acquisition of Dual or Triple Infection during Infancy. J Clin Microbiol 46, 507–514 (2008).

8. Bédarida, S., Dussol, B., Signoli, M. & Biagini, P. Analysis of Anelloviridae sequences characterized from serial human and animal biological samples. Infect Genetics Evol 53, 89–93 (2017).

9. Koonin, E. V., Dolja, V. V. & Krupovic, M. The healthy human virome: from virus–host symbiosis to disease. Curr Opin Virol 47, 86–94 (2021).

10. Tyschik, E. A., Rasskazova, A. S., Degtyareva, A. V., Rebrikov, D. V. & Sukhikh, G. T. Torque teno virus dynamics during the first year of life. Virol J 15, 96 (2018).

11. Virgin, H. W., Wherry, E. J. & Ahmed, R. Redefining Chronic Viral Infection. Cell 138, 30–50 (2009).

12. Cebriá-Mendoza, M. et al. Deep viral blood metagenomics reveals extensive anellovirus diversity in healthy humans. Sci Rep-uk 11, 6921 (2021).

13. Focosi, D., Antonelli, G., Pistello, M. & Maggi, F. Torquetenovirus: the human virome from bench to bedside. Clin Microbiol Infec 22, 589–593 (2016).

14. Arze, C. A. et al. Global genome analysis reveals a vast and dynamic anellovirus landscape within the human virome. Cell Host Microbe 29, 1305-1315.e6 (2021).

15. Calus, S. T., Ijaz, U. Z. & Pinto, A. J. NanoAmpli-Seq: a workflow for amplicon sequencing for mixed microbial communities on the nanopore sequencing platform. Gigascience 7, giy140. (2018).

16. Andrews, S. FastQC: A Quality Control Tool for High Throughput Sequence Data.

17. Ewels, P., Magnusson, M., Lundin, S. & Käller, M. MultiQC: summarize analysis results for multiple tools and samples in a single report. Bioinformatics 32, 3047–3048 (2016).

18. Bushnell, B. BBMap: A Fast, Accurate, Splice-Aware Aligner. https://www.osti.gov/biblio/1241166-bbmap-fast-accurate-splice-aware-aligner.

19. Wick, R. filtlong.

20. Wick, R. Porechop.

21. Baloğlu, B. et al. A workflow for accurate metabarcoding using nanopore MinION sequencing. Methods Ecol Evol 12, 794–804 (2021).

22. Sedlazeck, F. J., Rescheneder, P. & Haeseler, A. von. NextGenMap: fast and accurate read mapping in highly polymorphic genomes. Bioinformatics 29, 2790–2791 (2013).

23. Li, H. Aligning sequence reads, clone sequences and assembly contigs with BWA-MEM. (2013).

24. Li, H. & Durbin, R. Fast and accurate short read alignment with Burrows–Wheeler transform. Bioinformatics 25, 1754–1760 (2009).

25. Institute, B. Picard Tools.

26. Nurk, S., Meleshko, D., Korobeynikov, A. & Pevzner, P. A. metaSPAdes: a new versatile metagenomic assembler. Genome Res 27, 824–834 (2017).

27. Schmieder, R. & Edwards, R. Quality control and preprocessing of metagenomic datasets. Bioinformatics 27, 863–864 (2011).

28. Rognes, T., Flouri, T., Nichols, B., Quince, C. & Mahé, F. VSEARCH: a versatile open source tool for metagenomics. Peerj 4, e2584 (2016).

29. Nishimura, Y. et al. Environmental Viral Genomes Shed New Light on Virus-Host Interactions in the Ocean. Msphere 2, mSphere.00359-16, e00359--16 (2017).

30. Vaser, R., Sović, I., Nagarajan, N. & Šikić, M. Fast and accurate de novo genome assembly from long uncorrected reads. Genome Res 27, 737–746 (2017).

31. Camacho, C. et al. BLAST+: architecture and applications. Bmc Bioinformatics 10, 421 (2009).

32. Woodcroft, B. J., Boyd, J. A. & Tyson, G. W. OrfM: a fast open reading frame predictor for metagenomic data. Bioinformatics 32, 2702–2703 (2016).

33. Shen, W., Le, S., Li, Y. & Hu, F. SeqKit: A Cross-Platform and Ultrafast Toolkit for FASTA/Q File Manipulation. Plos One 11, e0163962 (2016).

34. Arze, C. A. et al. Global genome analysis reveals a vast and dynamic anellovirus landscape within the human virome. Cell Host Microbe 29, 1305-1315.e6 (2021).

35. Takahashi, K., Iwasa, Y., Hijikata, M. & Mishiro, S. Identification of a new human DNA virus (TTV-like mini virus, TLMV) intermediately related to TT virus and chicken anemia virus. Arch Virol 145, 979–993 (2000).

36. Bailey, T. L. et al. MEME SUITE: tools for motif discovery and searching. Nucleic Acids Res 37, W202–208 (2009).

37. Galmès, J. et al. Potential implication of new torque teno mini viruses in parapneumonic empyema in children. European Respir J 42, 470–479 (2013).

38. Tyschik, E. A., Shcherbakova, S. M., Ibragimov, R. R. & Rebrikov, D. V. Transplacental transmission of torque teno virus. Virol J 14, 92 (2017).

39. Maggi, F. et al. Dynamics of Persistent TT Virus Infection, as Determined in Patients Treated with Alpha Interferon for Concomitant Hepatitis C Virus Infection. J Virol 75, 11999–12004 (2001).

40. Mariscal, L. F. et al. TT Virus Replicates in Stimulated but Not in Nonstimulated Peripheral Blood Mononuclear Cells. Virology 301, 121–129 (2002).

41. Focosi, D., Macera, L., Boggi, U., Nelli, L. C. & Maggi, F. Short-term kinetics of torque teno virus viraemia after induction immunosuppression confirm T lymphocytes as the main replication-competent cells. J Gen Virol 96, 115–117 (2015).

42. Maggi, F. et al. TT virus (TTV) loads associated with different peripheral blood cell types and evidence for TTV replication in activated mononuclear cells. J Med Virol 64, 190–194 (2001).

43. Rosette-forming human lymphoid cell lines. I. Establishment and evidence for origin of thymus-derived lymphocytes - PubMed. https://pubmed.ncbi.nlm.nih.gov/4567231/.

44. Qiu, J. et al. Human circovirus TT virus genotype 6 expresses six proteins following transfection of a full-length clone. J Virol 79, 6505–6510 (2005).

45. Mushahwar, I. K. et al. Molecular and biophysical characterization of TT virus: Evidence for a new virus family infecting humans. Proc National Acad Sci 96, 3177–3182 (1999).

46. Okamoto, H. et al. Molecular cloning and characterization of a novel DNA virus (TTV) associated with posttransfusion hepatitis of unknown etiology1The nucleotide sequence data of the TTV isolate (TA278) reported in this paper will appear in the DDBJ, EMBL and GenBank nucleotide sequence databases with accession number AB008394.1. Hepatol Res 10, 1–16 (1998).

47. Todd, D., Creelan, J. L., Mackie, D. P., Rixon, F. & McNulty, M. S. Purification and biochemical characterization of chicken anaemia agent. J Gen Virol 71, 819–823 (1990).

48. Wang, X.-C. et al. Viral metagenomics reveals diverse anelloviruses in bone marrow specimens from hematologic patients. J Clin Virol 132, 104643 (2020).

49. Okamoto, H. et al. Circular Double-Stranded Forms of TT Virus DNA in the Liver. J Virol 74, 5161–5167 (2000).

50. Doan, T. et al. Paucibacterial Microbiome and Resident DNA Virome of the Healthy Conjunctiva. Invest Ophth Vis Sci 57, 5116–5126 (2016).

51. Smits, S. L. et al. High Prevalence of Anelloviruses in Vitreous Fluid of Children With Seasonal Hyperacute Panuveitis. J Infect Dis 205, 1877–1884 (2012).

52. Venkataraman, T. et al. Comprehensive profiling of antibody responses to the human anellome using programmable phage display. Biorxiv 2022.03.28.486145 (2022) doi:10.1101/2022.03.28.486145.

